# A Hypomorphic *Cystathionine ß-Synthase* Gene Contributes to Cavefish Eye Loss by Disrupting Optic Vasculature

**DOI:** 10.1101/805804

**Authors:** Li Ma, Aniket V. Gore, Daniel Castranova, Janet Shi, Mandy Ng, Kelly A. Tomins, Corine M. van der Weele, Brant M. Weinstein, William R. Jeffery

## Abstract

Vestigial structures are key indicators of evolutionary descent but the mechanisms underlying their development are poorly understood. This study examines vestigial eye formation in the teleost *Astyanax mexicanus*, which consists of a sighted surface-dwelling morph and different populations of blind cave morphs. Cavefish embryos initially develop optic primordia but vestigial eyes are formed during larval development. Multiple genetic factors are involved in cavefish eye loss but none of the mutated genes have been identified. Here we identify *cystathionine ß-synthase* (*cbsa*), which encodes the key enzyme of the transsulfuration pathway, as a mutated gene responsible for eye degeneration in multiple cavefish populations. The inactivation of *cbsa* affects eye development by inducing accumulation of the transsulfuration intermediate homocysteine and defects in optic vasculature, including aneurysms and eye hemorrhages, leading to oxygen deficiency. Our findings suggest that localized modifications in the circulatory system and hypoxia had important roles in the evolution of vestigial eyes in blind cavefish.

## Introduction

Vestigial structures, such as the reduced wings of flightless birds, the hind limb remnants of pythons, and the degenerate eyes of blind cavefish, were key factors in Darwin’s recognition of descent with modification during evolution. However, the genetic and physiological mechanisms responsible for the development of rudimentary structures are still poorly understood.

In the teleost *Astyanax mexicanus*, blind cavefish (CF) have been derived repeatedly from sighted surface fish (SF) ancestors^1–3^. The loss of eyes in *A. mexicanus* CF involves initial optic development followed by subsequent degeneration^1,2^. Eye primordia with a lens and retina are formed in CF embryos, but the lens undergoes massive apoptosis, the retina becomes apoptotic and disorganized, and eye growth is eventually arrested during larval development ^3–5^. Accordingly, the rate of optic growth fails to keep pace with the increase in overall body size as CF larvae develop into adults, and the small non-functional eyes are overgrown by skin and connective tissue. About 30 distinct CF populations with various degrees of eye reduction have evolved in Mexican caves after colonization by surface fish ancestors during the mid-to late Pleistocene, and several of these CF populations have evolved vestigial eye phenotypes independently ^6,7^. Similar eye reduction or loss occurs in other cave dwelling species^8^ and animals adapted to other perpetually dark habitats^9^.

The existence of hybridizable surface dwelling and cave adapted morphs in *A. mexicanus* has allowed eye degeneration to be studied by genetic methods^2^. These studies have shown that eye loss in the Pachón CF (PA-CF) population is controlled by multiple genetic factors, each controlling a part of the complex vestigial eye phenotype^10–12^. Furthermore, complementation crosses between different CF populations have demonstrated that some of the genetic factors involved in eye loss are the same whereas others are unique^13^. In addition to genetic changes, epigenetic events may also have roles in the evolution of CF eye loss^14^. Quantitative trait locus (QTL) analysis revealed about 15 non-overlapping genomic regions that are responsible for lens and eye reduction in PA-CF^10–12,15^. The alignment of eye QTL with the PA-CF genome sequence has suggested numerous candidates for genes controlling vestigial eye formation,^15^. However, the identities of the inactivated genes within these QTL intervals, the type of mutations that are present, and the genetic mechanisms operating to generate the eyeless phenotype, have not been established.

Here we identify *cystathionine ß-synthase a* (*cbsa*) as one of the inactivated genes responsible for eye degeneration in multiple *A. mexicanus* CF populations. Studies of the hypomorphic *cbsa* phenotype revealed a novel mechanism for arresting eye growth based on accumulation of the transsulfuration intermediate homocysteine and disruption of optic vasculature. Our findings suggest that interference with normal circulatory system function by a *cbsa* mutation may have played a critical role in the evolution of CF eye degeneration.

## Results

### *cbsa* as a candidate gene for cavefish eye loss

To identify a mutated gene responsible for CF eye loss, we examined the *A*. *mexicanus* Ensembl Scaffold KB871589.1 from, which contains the peak marker of an eye size QTL located between *hsp2bp* and *ankrd10a* (Fig. 1a)^11–13,15^. According to the Ensembl AstMex 1.0.2 genome assembly, this scaffold contains 21 predicted protein-coding genes. We surveyed these genes by qualitative RT-PCR for expression differences in SF and PA-CF at 40 hpf (Fig. 1a; Table S1), when many changes associated with eye degeneration first appear in CF^1–5^. Since CF eye degeneration is a recessive trait^2–5^, we focused on genes that are downregulated in PA-CF relative to SF, and four such genes were identified: *cystathionine ß-synthase a* (*cbsa*), *αA-crystallin* (*cryaa*), *heat shock factor 2 binding protein* (*hsf2bp*), and *cysteinyl-tRNA synthetase 2* (*cars2*) (Fig. 1a). We next focused on protein coding genes expressed in the eyes. Accordingly, the *cars* gene was not pursued further because it encodes a mitochondrial aminoacyl-tRNA synthetase^16^. Of the remaining genes, the *cryaa* gene, although expressed in the lens, was also not studied further because it was previously shown to be under *trans*-acting regulation during CF development^17^. The *hsf2bp* gene encodes a heat shock transcription factor 2 binding protein^18^. We used *in situ* hybridization to determine the pattern of *hsf2bp* expression during normal culture at 25°C or following a 1 hour 37°C heat shock (Fig. 1b). The results indicated that *hsf2bp* expression was ubiquitous in SF and PA-CF, and although expression was increased during a heat shock, the increase was not particularly strong in the eyes (Fig. 1b), so we also excluded this gene from further consideration. Thus, our attention ultimately focused on *cbsa*, a gene that is expressed in the developing eyes and other locations in *A. mexicanus* (see below) and was previously predicted as a candidate for congenital anomaly in the CF eye loss by Ingenuity Pathway Analysis of gene expression data^15^.

**Fig. 1:**
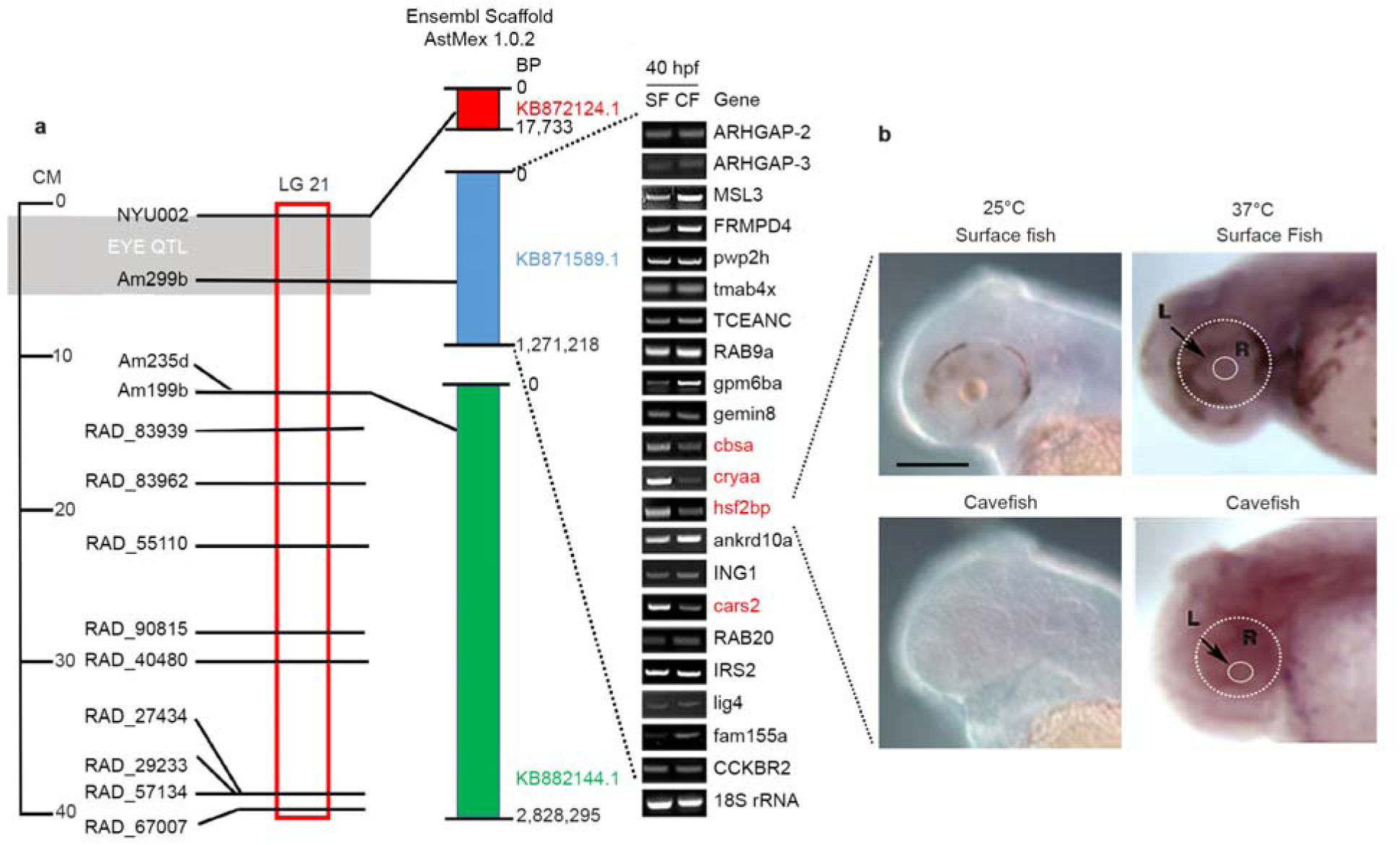
Screen of PA genomic scaffold KB871589.1 containing a peak eye QTL for downregulated genes expressed in SF eyes at 40 hpf. (**a**) Location of an eye QTL on *A. mexicanus* linkage group 21 (left), alignment to nearby scaffolds of the draft PA-CF (AstMex1.0.2) genome sequence based on the position of markers (center), and semi-quantitative RT-PCR analysis of gene expression of 21 predicted scaffold genes at 40 hpf (right). Expression of genes in SF and CF determine by qualitative RT-PCR are shown from top to bottom according to their 5’ to 3’ order on the scaffold. CM: Centimorgans. LG: linkage group. See Table S1 for PA-CF gene IDs. Genes downregulated in CF compared to SF are shown in red. (**b**) *In situ* hybridization showing *hsf2bp* expression in 40 hpf SF and PA-CF embryos at 25°C (left) and after treatment at 37°C for 1 hour (right). Dashed lines encircle the eyes and lenses on the right frames. L: lens. R: retina. Scale bar: 150 μm; same magnification in all frames.

We first examined the expression pattern of *cbsa* and its paralog *cbsb* during SF and PA-CF development by *in situ* hybridization with gene specific probes (Fig. 2a-p). At early stages of embryonic development (10–20 hours), the patterns of *cbsa* and *cbsb* expression were similar in SF and PA-CF, including in the developing eye primordia, although *cbsa* expression was markedly weaker in PA-CF than in SF (Fig. 2a–f), and *cbsb* expression was generally stronger than *cbsa* in PA-CF (Fig. 2a–f and i–n). By 40 hours of development, *cbsa* and *cbsb* expression was seen in many developing organs (Fig. 2g, h, o and p). The expression of *cbsa* was strong in the SF head, including the eyes and parts of the brain, heart, pectoral fin buds, pancreas, liver, and myotomes (Fig. 2h). The pattern of *cbs* gene expression in *Astyanax* surface fish closely resembled that reported in zebrafish^19^. In PA-CF *cbsa* expression was downregulated in all of these organs except the pancreas (data not shown) and liver (Fig. 2h). The expression of *cbsb* was similar in most of these organs in SF and PA-CF at 40 hpf, with the notable exception of the head and myotomes, where *cbsb* expression was reduced in both PA-CF and SF (Fig. 2o and p). The results suggest that *cbsa* and *cbsb* show mostly overlapping expression in during SF and PA-CF development, except for the PA-CF eyes and myotomes in which neither of the *cbs* paralogs was strongly expressed. The results suggested that *cbsb* expression could compensate for downregulated *cbsa* expression is all parts of CF larvae except for the eyes. The *in situ* hybridization results were confirmed by comparing *cbsa* and *cbsb* expression in isolated larval heads and trunks by qualitative and quantitative RT-PCR at 40 hpf, which showed that *cbsa*, but not *cbsb*, is significantly downregulated in PA-CF larval heads at this developmental stage (Fig. 2q).

**Fig. 2:**
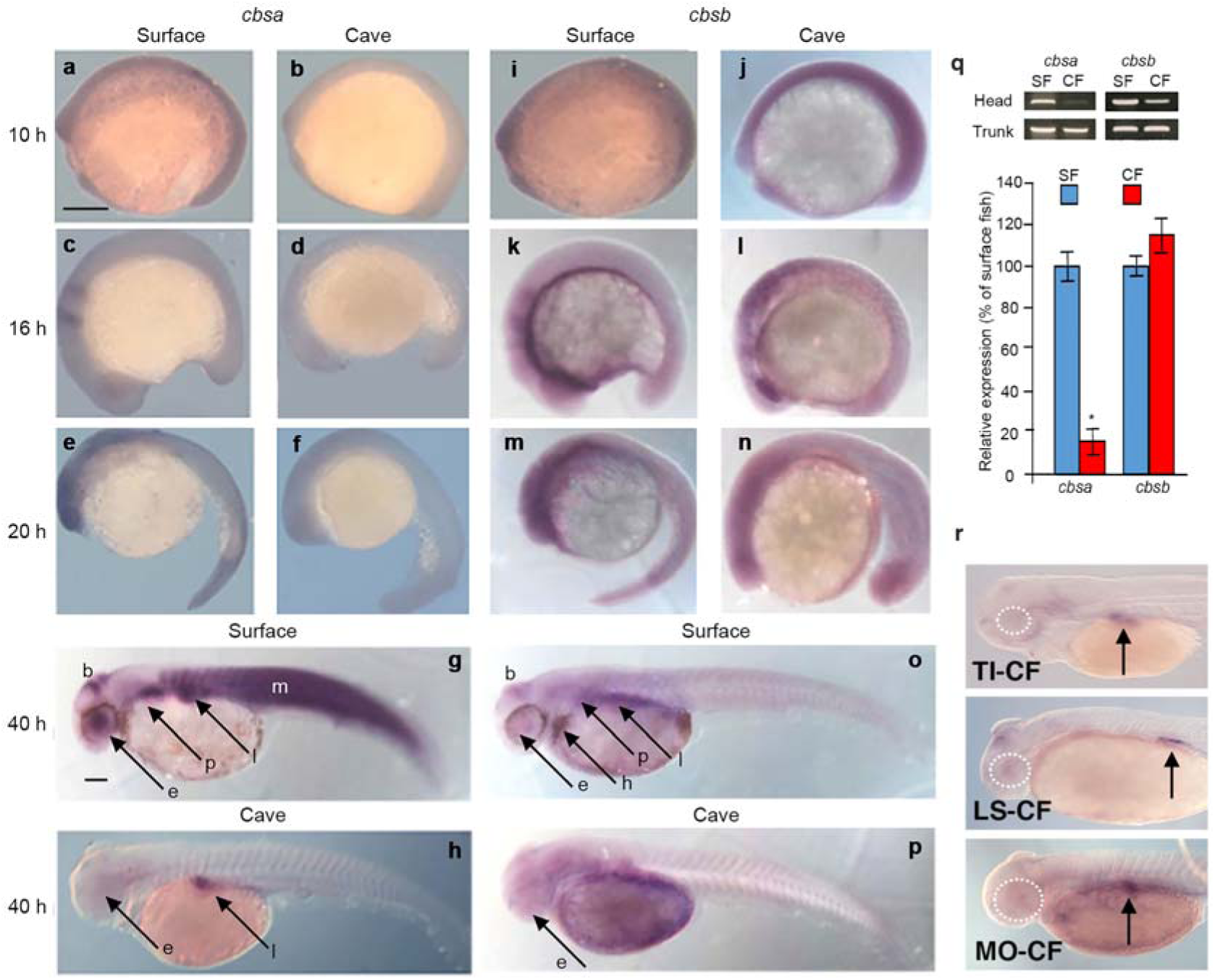
Expression of *cbsa* and *cbsb* in surface fish and cavefish. (**a-p**). *In situ* hybridization showing *cbsa* (**a-h**) and *cbsb* (**i-p**) expression during SF and PA-CF development. b: brain. e: eyes. h: heart. p: pectoral fin buds. l: liver. M: myotomes. **q**. Qualitative (top) and quantitative RT-PCR comparing the levels of cbsa and cbsb expression in cavefish heads and trunks and heads, respectively. Histograms show means, and error bars represent s. e. m. of three biological replicates. Asterisk: P < 0.05. Significance was determined by a two-tailed Students t test. **r**. *In situ* hybridization showing cbsa expression in TI-CF, SF-CF, and MO-CF at 40 hpf. Arrows: liver. Scale bar in A is 200 μm; magnification is the same in a–n and i-n. Scale bar in g is 300 μm; magnification is the same in g, h, o, p and r.

The results described above suggested that *cbsa* expression is downregulated in the anterior regions of PA-CF larvae, including the eyes. Eye loss is thought to have evolved independently at least twice in different *A. mexicanus* CF populations^1,2,13^. Therefore, we conducted *in situ* hybridization to determine whether *cbsa* expression is also changed in additional CF populations. The results showed similar patterns of *cbsa* downregulation at 40 hpf, including the degenerating eyes, brain, and myotomes, in PA-CF, TI-CF, LS-CF, and MO-CF (Fig. 2r). The results establish *cbsa* as a candidate for controlling eye loss in multiple CF populations.

### The cavefish *cbsa* gene contains a hypomorphic mutation

We next asked whether downregulation of the *cbsa* gene in PA-CF is caused by *cis*- or *trans*-acting regulation by conducting F1 hybrid tests^15,20^. In these experiments, SF females were crossed with PA-CF males to generate F1 hybrids, RNA was extracted from the heads of the F1 hybrid and parental larvae at 40 hpf, and a portion of the *cbsa* coding region containing an SNP marker distinguishing the SF and CF alleles was amplified by PCR, cloned, and sequenced (Fig. 3a). The results showed that expression of the PA-CF *cbsa* allele was not increased in the F1 hybrid background, as would be expected under *trans*-acting control, thus implicating *cis*-regulation as the cause of *cbsa* downregulation.

**Fig. 3:**
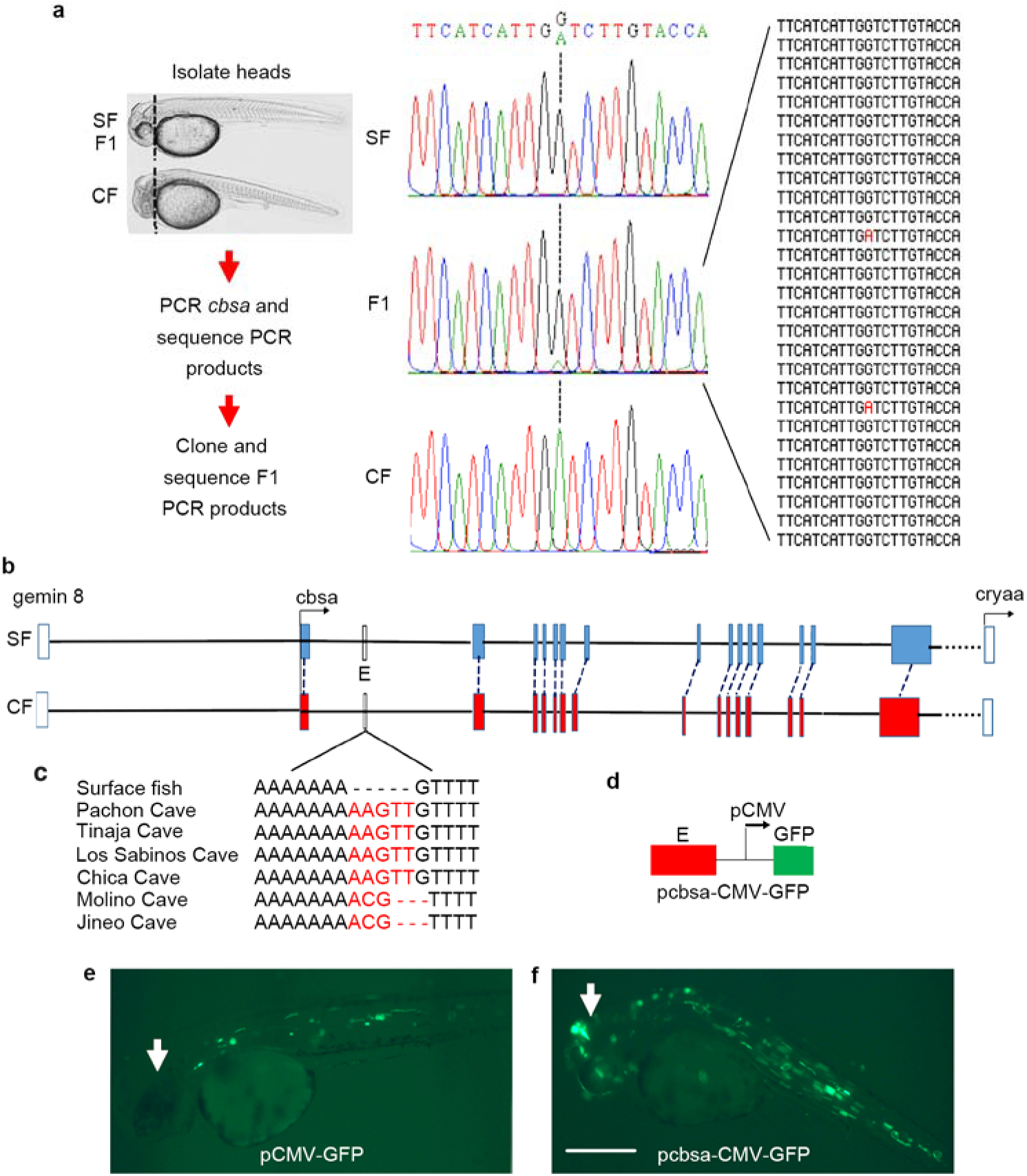
*cbsa* gene regulation and structure. (**a**) F1 hybrid test. Left: Summary of procedure. Middle: Sequenced *cbsa* RT-PCR products from SF, PA-CF, and F1 hybrid heads showing SF and CF *cbsa* marker SNP (A and G nucleotides, respectively). Right: Marker region sequences of TOPO cloned RT-PCR products from F1 hybrid heads showing the CF allele marker in red. (**b, c**) Maps of the SF and PA-CF *cbsa* loci showing indels in CF populations (c, red letters) and predicted enhancer (E) in SF *cbsa*. Colored boxes: exons. White boxes: Flanking *gemin8* and *cryaa* loci. (**d-f**) Expression of CMV-GFP constructs at 40 hpf. (d) *cbsa* enhancer E-CMV-GFP construct. (e) Control pCMV-GFP did not drive GFP expression in the head (arrow) in 54 injected SF embryos. (e) pcbsa-enhancer-CMV-GFP drove GFP expression in the head (arrow), including the eyes and brain, in 75 of 101 injected SF embryos.

To identify potential *cis*-regulating mutations, we sequenced the SF and PA-CF *cbsa* loci and immediate flanking regions. Although the draft genome sequences of SF and PA-CF are available in Ensembl (Astyanax_mexicanus-2.0 and AstMex1.0.2 respectively), gaps and possible assembly problems did not permit reliable detection of putative mutations across the large genomic span of the *cbsa* gene locus. Therefore, we re-sequenced about 32 kb of PA-CF and SF genomic DNA spanning the *cbsa* loci and flanking non-coding regions between the upstream *gemin8* and downstream *cryaa* genes. The results showed that the SF and CF *cbsa* genes consist of 15 exons (Fig. 3b). We used RACE to identify the SF and CF *cbsa* transcripts and to survey them for possible sequence differences. The 5’ non-coding regions of CF and SF *cbsa* mRNAs, which correspond to the first non-coding exon and the first few bases of the second exon, showed only two nucleotide differences. In addition, the CF *cbsa* coding region has no internal stop codons, suggesting that it encodes a complete CBSA protein (Fig. S1a). The deduced PA-CF and SF CBSA proteins are 99% identical, differing by only five amino acids (Fig. S1b). Only one of the divergent amino acids (G 257) is conserved in other known vertebrate CBS proteins. Although it is possible that PA-CF CBSA is functionally different from SF CBSA because it is missing a run of two amino acids (G257-K258; Fig. S1c) that are present in other teleost CBS proteins, this difference cannot account for hypomorphic activity of the PA-CF *cbsa* gene.

Outside the protein coding regions, there are only a few substitutions in the 7 kb non-coding region located between *cbsa* and *gemin8*, which shows 99% nucleotide identity between PA-CF and SF, or in the 5 kb non-coding region located between *cbsa* and *cryaa*, which is also highly conserved in SF and PA-CF^17^. In addition, *cbsa* introns are also highly conserved in PA-CF, except for the following: a 5 bp insert in intron 1, a 323 bp deletion in intron 6, and a 120 bp insertion in intron 8 (Table S1). To explore the possibility that one or more of these PA-CF *cbsa* introns contain potential *cis*-regulatory changes, we amplified and compared the sequences surrounding the changes in introns 1, 6, and 8 in the PA-CF, TI-CF, LS-CF, CH-CF, JI-CF, and MO-CF *cbsa* genes. Genetic studies have shown that eye loss evolved independently at least twice among some of these CF populations^2,11,13^, and our *in situ* hybridization data indicated that *cbsa* is downregulated in PA-CF, TI-CF, LS-CF, and MO-CF eyes (Fig. 2). We found that the indels in introns 6 and 8 were present in some, but not all, of these CF populations (Table S1), suggesting that they are unlikely to be generally responsible for *cbsa* downregulation. In contrast, small inserts consisting of two different sequences were found at the same position in intron 1 in all six CF populations (Fig. 3c), suggesting that this region may include a *cis*-acting mutation.

We next searched for enhancers in the genomic region between the 3’ termini of the *gemin8* and *cbsa* genes using iEnhancer-2L^21^. This tool predicted 5 putative enhancers, but most of them showed little or no sequence differences between PA-CF and SF, and no enhancers were predicted within *cbsa* introns 6 or 8. However, a putative enhancer (E region, Fig. 1b) was detected in intron 1 of *cbsa*, which includes the indels in all six CF populations. When a SF 1040 bp DNA fragment including the E region was inserted into the pCMV-GFP expression vector and injected into SF eggs, the reporter construct drove GFP expression in the myotomes, brain, and eyes of 40 hpf SF embryos (Fig. 3d-f). These results support the presence of a *cis*-acting mutation in an enhancer located in the large first intron of the PA-CF *cbsa* gene, although other *cis*-acting changes, within or beyond the currently sequenced regions, may also contribute to *cbsa* downregulation during PA-CF development. These results (and functional data below) suggest that *cbsa* could be one of the mutated genes responsible for CF eye loss.

### Cavefish show homocystinuria-like features

The *cbsa* gene encodes cystathionine ß-synthase (CBS), the limiting enzyme of the methionine/transsulfuration pathway, which converts homocysteine (hCys) to cystathionine, a precursor of cysteine and glutathione^22^ (Fig. 4a). Mutations in the human *cbs* gene are the major cause of homocystinuria, a disorder in methionine metabolism leading to toxic hCys accumulation and eye abnormalities^23,24^. In fact, the small eyes of CF larvae resemble the eyes of some homocystinuria patients in showing an “off-center” lens^2–5^. To test the possibility that CF show homocystinuria-like features, we compared hCys levels by ELISA quantification and determination of the expression of *activating transcription factor 3* (*atf3*), a major hCys-sensitive gene^25^, in CF and SF larvae. There ELISA results showed 4–6 fold increases of hCys levels in PA-CF and TI-CF larvae at 40 hpf (Fig. 4b). We also detected strong upregulation of the *atf3* gene in PA-CF and TI-CF relative to SF between about 2 and 4 days post-fertilization (dpf) (Fig. 4c). To determine the developmental effects of elevated hCys, we injected hCys into SF eggs and examined larval morphology. The results showed that about 21% of the SF larvae developing from hCys injected eggs showed reduced eye size, the displacement of the lens to the ventral side of the eye, or complete eye loss, although other morphological features were not visibly affected (Fig. 4d, f, and g; Table S2a). In contrast, much fewer control larvae developing from eggs injected with cysteine showed effects on eye development (Fig. 4a, e; Table S2a). Together, these results indicate that CF exhibit homocystinuria-like features that impact eye development.

**Fig. 4:**
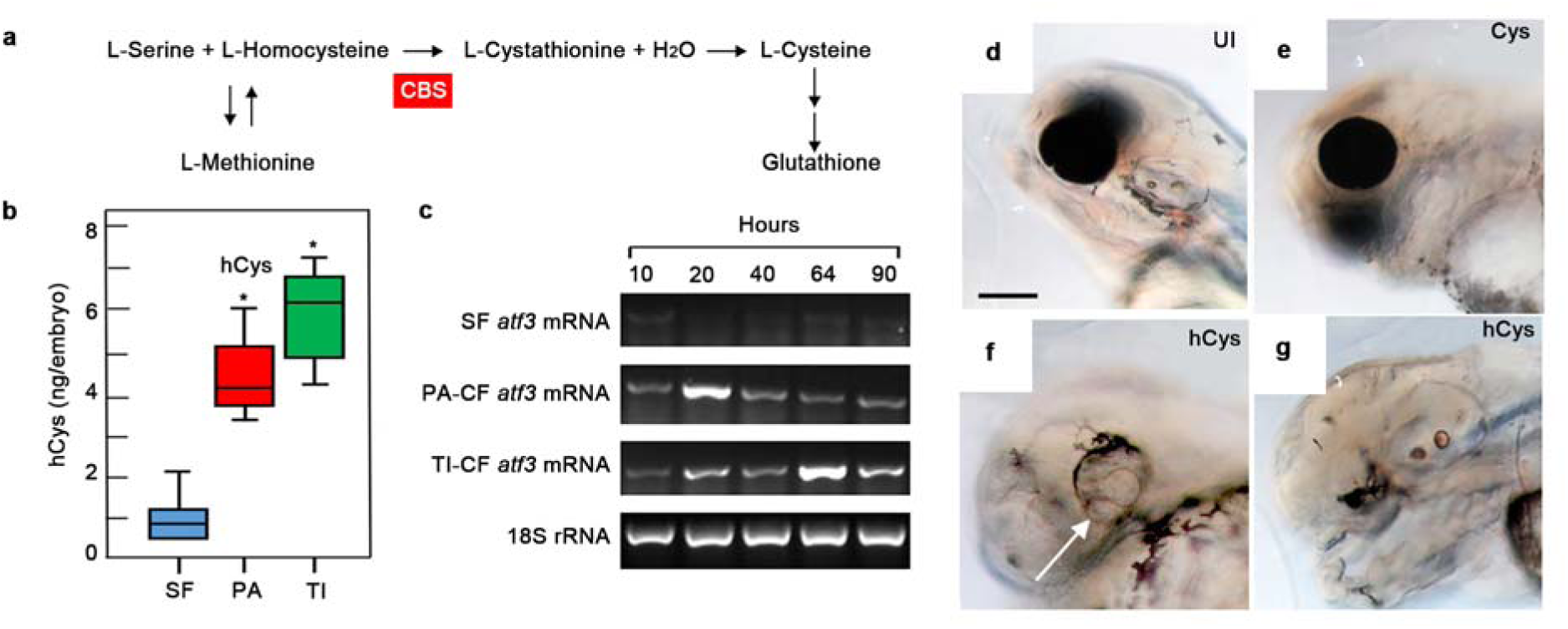
Homocysteine elevation in cavefish and effects on surface fish eye development. (**a**) The transsulfuration pathway showing one of the reactions catalyzed by the CBS enzyme. (**b**) ELISA determination of hCys levels in PA-CF and TI-CF relative to SF at 40 hpf. Box plots show medians, Q1 and Q3, and errors bars represent the whiskers of six biological replicates. Asterisks: *P* < 0.001. Significance was determined by two-tailed Students *t* test. *n* =14 for each box plot. (**c**) Expression of *atf3* during PA-CF, TI-CF, and SF development determined by qualitative RT-PCR. (**d-g**) Effects on eye development at 4 dpf after injection of 5 mM hCys into SF eggs (f). (d, e) Normal eye development in un-injected (UI) SF eggs (d) and SF eggs injected with 5 mM cysteine (Cys) (e). (f, g) Development of a small eye with ventrally displaced lens (arrow, F) or complete suppression of eye formation (g) in SF eggs injected with 5 mM hCys. Scale bar: 150 μm; same magnification in D to F.

### Defects in cavefish optic vasculature

High hCys levels have effects on cardiovascular function, increasing the risk of blood clots, aneurysms, and hemorrhagic stroke in homocystinuria patients and *Cbs* +/− mice^26,27^. Therefore, we examined vasculature development and function in CF by microangiography, blood cell staining, and eye imaging. The results revealed leaky optic and brain vasculature, erythrocyte pooling and aneurysms around the eyes, and eye hemorrhages in PA-CF larvae, but not in SF larvae (Fig. 5a-i). Eye hemorrhages were frequently observed in CF larvae (Fig. 5g–h, j), but not in more than 1000 SF larvae evaluated in this study (Fig. 5f). The eye hemorrhages were generally unilateral, ranging from small foci on the dorsal side of the eye to large areas of the orbit, and were present for only about 3–4 days in most larvae (Fig. 5l). During this period, the degenerating CF eyes were invaded by phagocytic macrophages, probably to remove leaked blood cells (Fig. 5k). Survival studies showed that PA-CF larvae with eye hemorrhages have similar viability to SF larvae (Fig. 4m), showing that defective optic vasculature does not decrease CF viability. The results indicate that optic vasculature is defective in CF.

**Fig. 5:**
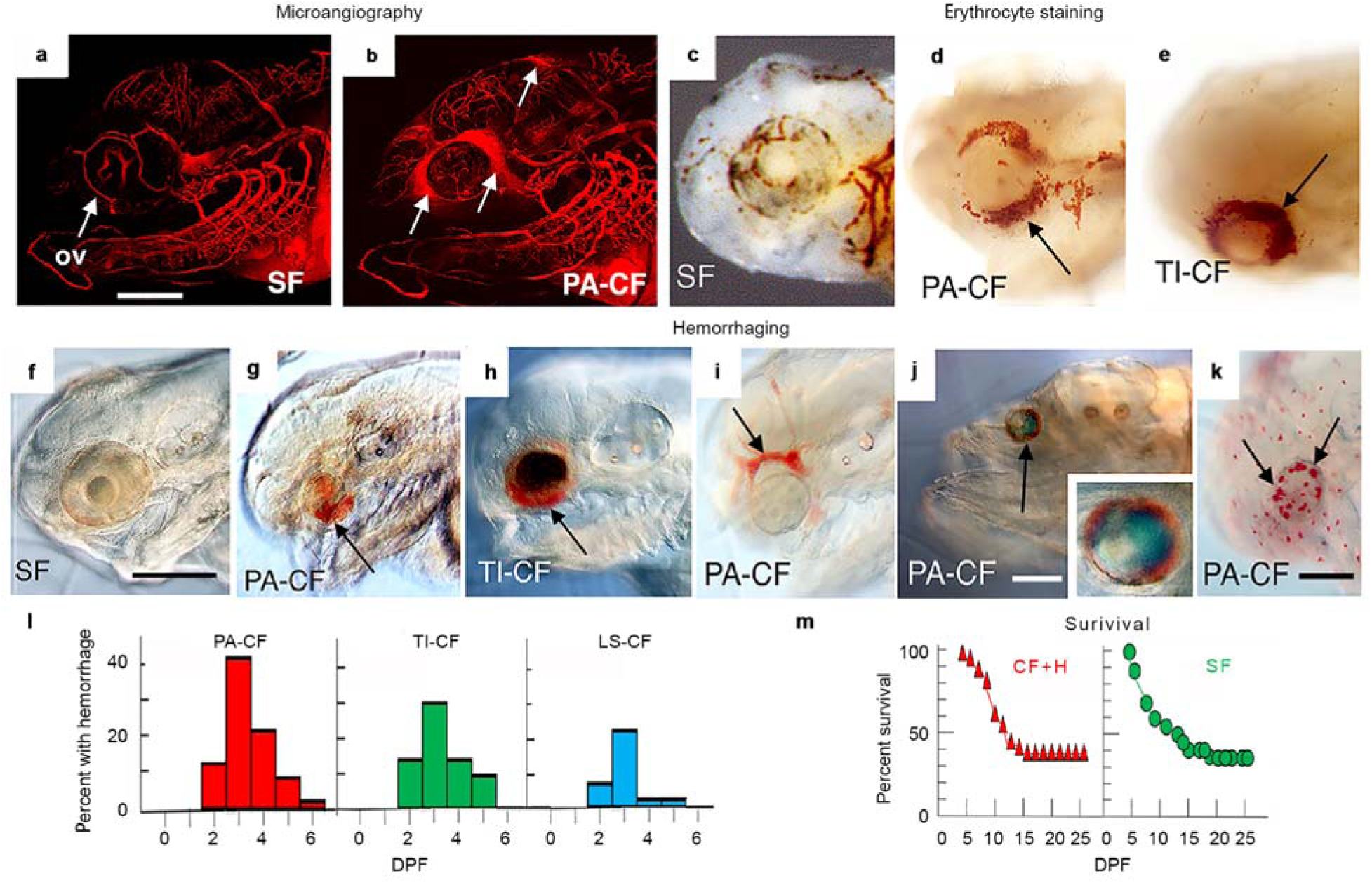
Defective optic vasculature in cavefish. (**a** and **b**) SF (a) and PA-CF (b) at 5 days post-fertilization (dpf) showing leaky vasculature around the PA-CF eye. ov: optic vasculature. Each assay was conducted with three biological replicates. (**c** to **e**) SF (c), PA-CF (d), and TI-CF (e) showing pooled erythrocytes (arrows in d, e) around CF eyes at 2.5 dpf. n = 24 for SF and 36 for PA-CF. (**f** to **h**) SF (f), PA-CF (g), and TI-CF (h) at 4 dpf. showing hemorrhages (arrows in G, H) in CF eyes (**i)**. Aneurysm (arrow) in PA-CF optic vasculature at 4 dpf. (**j**) PA-CF showing eye hemorrhage (arrow) at 6 dpf (inset, 2.5 magnification of eye area). (**k**) Neutral red staining of PA-CF at 7 dpf showing macrophages (arrows) in degenerating eyes (36 of 36 specimens). Scale bars: (a) 150 μm (same magnification in a and b), (f) 100 μm (same magnification in C–I), (j and k) 150 μm. (**l**) The percentage of PA-CF (n = 221), TI-CF (n = 125), and LS-CF (n = 86) with hemorrhages during larval development. (**m**) Survival of SF (right) (n = 132) and PA-CF (n = 165) with hemorrhages (left) during larval development.

### Role of *cbs* genes in eye and optic vasculature development

To address the role of the *cbs* genes in eye and vasculature development, we knocked down *cbsa* and *cbsb* expression by injecting SF eggs with splice-inhibiting morpholinos (MO) (Fig. 6a-c). To validate this approach, we amplified and sequenced PCR products from the *cbs* morphants and obtained products of two different sizes: one with sequence corresponding to the processed *cbsa* exon 5 and 6 or *cbsb* exon 3 and 4, and the other with sequence corresponding to unprocessed introns containing the expected translation stop sites (Fig. S2a-d). The *cbsa* and *cbsb* MOs were specific for the corresponding genes respectively: PCR products derived from unprocessed mRNAs were obtained only when *cbsa* or *cbsb* MOs were injected, and only products corresponding to processed mRNAs were obtained when control MO was injected (Fig. S2a, b). The specific effects of the *cbs* MOs on eye development were also confirmed by controls showing that expression of the *major intrinsic protein* (*mip*), *cryaa*, *orthodenticle homeobox2* (*otx2*), and *SRY (sex determining region Y)-box2* (*sox2*), all of which are expressed in developing *Astyanax* forebrains and eyes^14,15^, decreased in the eyes (when present; see below) of *cbsa* and *cbsb* morphants, but not in control morphants (Fig. S2g, h). In contrast, there were no differences in *otx2* or *sox2* expression between the forebrains of control and *cbs* morphants (Fig. S2h). The results demonstrate specificity of the respective MOs for knocking down the *cbsa* or *cbsb* genes.

**Fig. 6:**
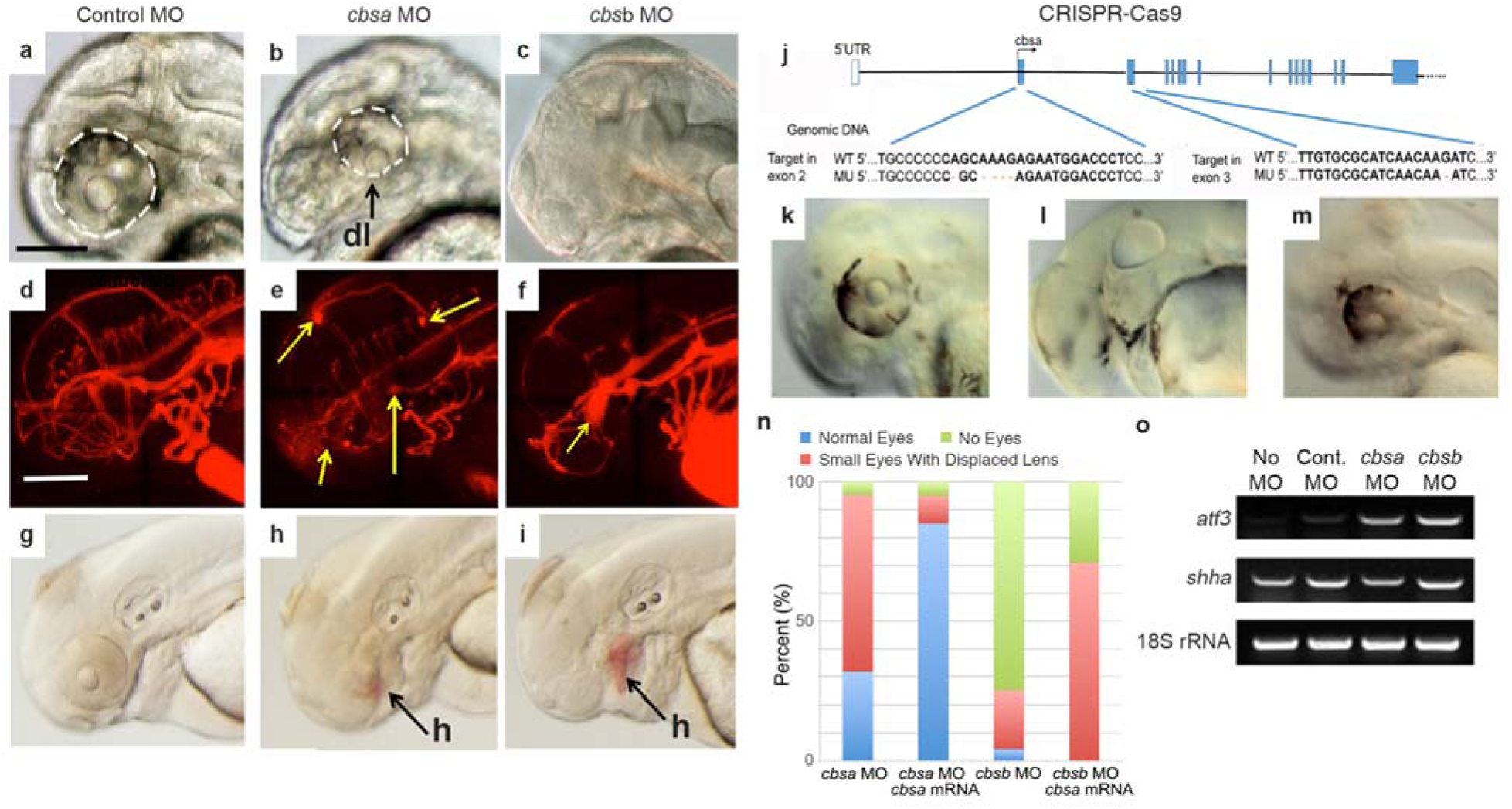
Effects of inhibiting *cbs* gene knockdown on eye and optic vasculature development. (**a** to **j**) Effects of splice-inhibiting morpholino (MO) knockdown of *cbsa* and *cbsb* genes (a-i) Effects of control (a, d, g), *cbsa* (b, e, h), and *cbsb* (c, f, i) MOs on eye development (a to c), optic vasculature integrity determined by microangiography (d-f), and optic hemorrhage formation (g-i) in SF at 40 hpf. Each assay was conducted with three biological replicates. Dashed lines: eyes. Yellow arrows: leaky vasculature. dl: displaced lens. h: hemorrhages. (j-m) Effects of CRISPR-Cas9 editing of cbsa on eye development in SF at 40 hpf. (j) Mutations induced in cbsa exons 3 and 4. k-m. Normal eyes in controls (k), missing eyes in exon 3 mutated larvae (l), and small eyes with ventrally displaced lenses in exon 4 mutated larvae. (**n**) Rescue of *cbsa* and *cbsb* MO induced eye defects in SF morphants by injection of *cbsa* mRNA. Bar graphs show percentage of normal and abnormal eye phenotypes. See Supplementary Table 2c for numbers. Effects of *cbs* MO knockdown on *atf3* gene expression, used as a proxy for homocysteine accumulation, and control *shha* gene expression in SF at 40 hpf.

We next determined the effects of *cbs* MO knockdown on eye development. The *cbsa* morphants developed a large number of small eyes with ventrally-displaced lenses, the typical CF eye phenotype^2–5^, and most of the *cbsb* morphants showed small eyes or completely missing eyes (Fig. 6b and c, n; Fig. S2g; Table S2b, c). No other gross morphological deficiencies were noted in the *cbs* morphants (Fig. S2g). Control morphants did not show these optic phenotypes (Fig. 6a; Fig. S2g; Table S2b, c). Co-injection of *cbsa* and *cbsb* MOs generated the same eye phenotypes as single MO injections, suggesting functional redundancy of the paralogous *cbs* genes (data not shown). The effects of *cbsa* MOs and *cbsb* MOs on eye development could be rescued by co-injection of *cbsa* mRNA (Fig. 6n), confirming the specificity of the *cbs* MO effects.

The effects of *cbsa* MO knockdown on SF eye development were confirmed by CRISPR-Cas9 editing of the *cbsa* gene. Using this method, mutations were introduced into exons 2 or 3 of *cbsa* (Fig. 6j) with about 29% efficiency (N = ???). Similar to the *cbs* morphants described above, larvae with CRISPR-Cas9 mutations showed abnormal or missing eyes (Fig. 6k-m). Together, the MO knockdown and CRISPR-Cas9 editing results indicate that expression of the *cbs genes* is required for normal eye development in *Astyanax*.

We also examined the roles of *cbs* MO knockdown on hCys accumulation and optic vasculature development. As a proxy for hCys levels, we determined *aft3* expression at 40 hpf in *cbsa* and *cbsb* morphants and found markedly increased expression of *atf3*, but not the control gene *sonic hedgehog a* (*shha*), in both cases (Fig. 6o). These results are consistent with the conclusion that *cbs* knockdown causes an increase in hCys levels. To determine the effects on vasculature development, we conducted microangiography in *cbs* and control MO-injected larvae. The *cbsa* morphants showed leaky vasculature around the developing eyes and the brain, and *cbsb* morphants showed leaky optic vasculature (Fig. 6d-f), or failed to develop cranial vasculature (data not shown), whereas vascular integrity was not compromised in control morphants (Fig. 6d). In addition, imaging indicated that some *cbsa* and *cbsb* morphants developed visible hemorrhages around the eyes and other cranial areas (Fig. 6h and i), but no hemorrhages were observed in control morphants (Fig. 6g). These results demonstrate that elevated hCys and optic vasculature phenotypes similar to those of CF can be generated by *cbs* gene knockdown in SF.

In summary, the results show that blocking *cbs* gene expression can promote higher levels of hCys, induce defective optic vasculature, and generate abnormal eye morphology, implying that disruption of optic vasculature may be responsible for CF eye loss.

### Inhibition of circulatory function affects eye growth in *Astyanax*

To further understand the role of the circulatory system in CF eye degeneration, we knocked down expression of the *rap1b* gene in SF. The *rap1b* gene disrupts endothelial junctions and is critical for normal vasculature development in zebrafish^28^. The *rap1b* morphants lacked erythrocytes in and around the eyes, implying that normal vascularization of the eye was defective (Fig. 7a). The *rap1b* morphants and controls initially showed eyes of similar size (Fig. 7b, d). Later in development, however, the *rap1b* morphants exhibited significantly smaller eyes than controls (Fig. 7c, e), demonstrating that knockdown *rap1b* expression affects eye growth in *Astyanax*. These results suggest that the effects of the hypomorphic *cbsa* gene on eye growth can be phenocopied by blocking blood circulation in *rap1b* morphants and provide evidence that defective optic vasculature may be an important factor in CF eye degeneration.

**Fig. 7.**
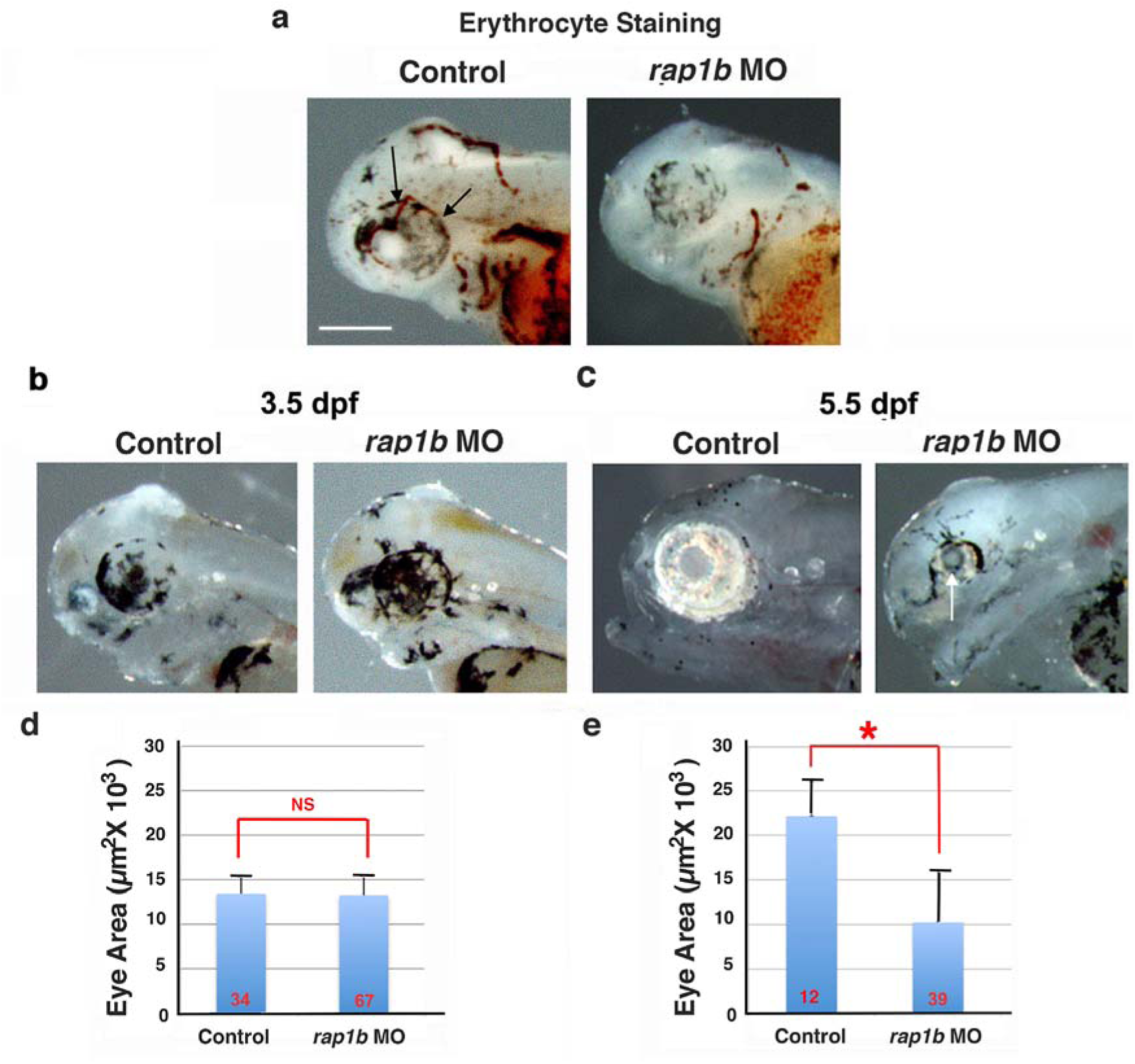
Effects of *rap1b* gene knockdown on circulatory system and eye development. (**a**) Erythrocytes stained by o-dianisidine at 2.5 dpf in un-injected control and *rap1b* morphants. Arrows indicate red blood cells. Scale bar: 300 μm same in a-c. (**b** and **c**) Eye morphology at 3.5 and 5.5 dpf in un-injected control and *rap1b* morphant larvae. Arrow indicates smaller eye in *rap1b* morphant at 5.5 dpf. (**d** and **e**). Bar graphs comparing mean eye size in control and *rap1b* morphant larvae at 3.5 dpf (**b**) and 5.5 dpf (**e**). Number of larvae analyzed are shown at the base of each bar. Error bars: SEM. NS indicates no significance and asterisk indicates significance at *P* < 0.001 in a Students two-tailed *t* test.

### Degenerating Cavefish Eyes Exhibit Hypoxia

One of the functions of optic vasculature is to supply the growing larval eyes with oxygen. Therefore, the discovery of defective optic vasculature opened the possibility that CF eyes may be deficient in oxygen. To explore this possibility, the *Astyanax* homologs of genes that that are highly sensitive to hypoxia in zebrafish^29,30^ were identified and their expression levels compared in isolated SF and PA-CF heads and trunks at 36 hpf (Fig. 8a, b). The hypoxia-sensitive genes examined were *hypoxia inducible factor 1*α*a* (*hif1*α*a*) and *hif1*α*b* (*hif1*α*b*), which encode master regulators of hypoxia^31^, *hemopexin* (*hpx*), a major hemolysis-sensitive heme scavenger^32^, *myoglobin* (*mb*), an oxygen transporter in muscle cells^33^, and *oxidative stress growth inhibitor 1* (*osgn1*). The *fatty acid desaturase 2* (*fads2*) and *ribosomal protein s3a* (*rps3a*) genes were selected as internal references. The results revealed that the *hif1*α*a*, *hpx*, and *mb* genes were significantly upregulated in the heads of PA-CF compared to SF and to PA-CF trunks (Fig. 8a, b). The *fads2* gene, which is downregulated during hypoxia in zebrafish^29^, was slightly increased in PA-CF heads compared to trunks, but the *rps3a* gene, which is upregulated during hypoxia in the liver of another teleost^34^, was also up-regulated in PA-CF relative to SF larval trunks (which contain the developing liver), although it was expressed at about the same level in heads (Fig. 8a, b). We also used antibody staining to determine the expression of the master hypoxia regulating factor HIF1α during SF and CF eye development. Early in development, when both SF and PA-CF eyes are both increasing in size, HIF1α was not detected in SF or PA-CF eyes, but as optic development proceeded and growth was arrested, PA-CF but not SF eyes showed HIF1α antibody staining, which continued to be present throughout the subsequent period of eye degeneration in (Fig. 8c-e). Together, the results of the hypoxia gene expression studies and HIF1α antibody staining suggest that degenerating PA-CF eyes exhibit hypoxia.

**Fig. 8.**
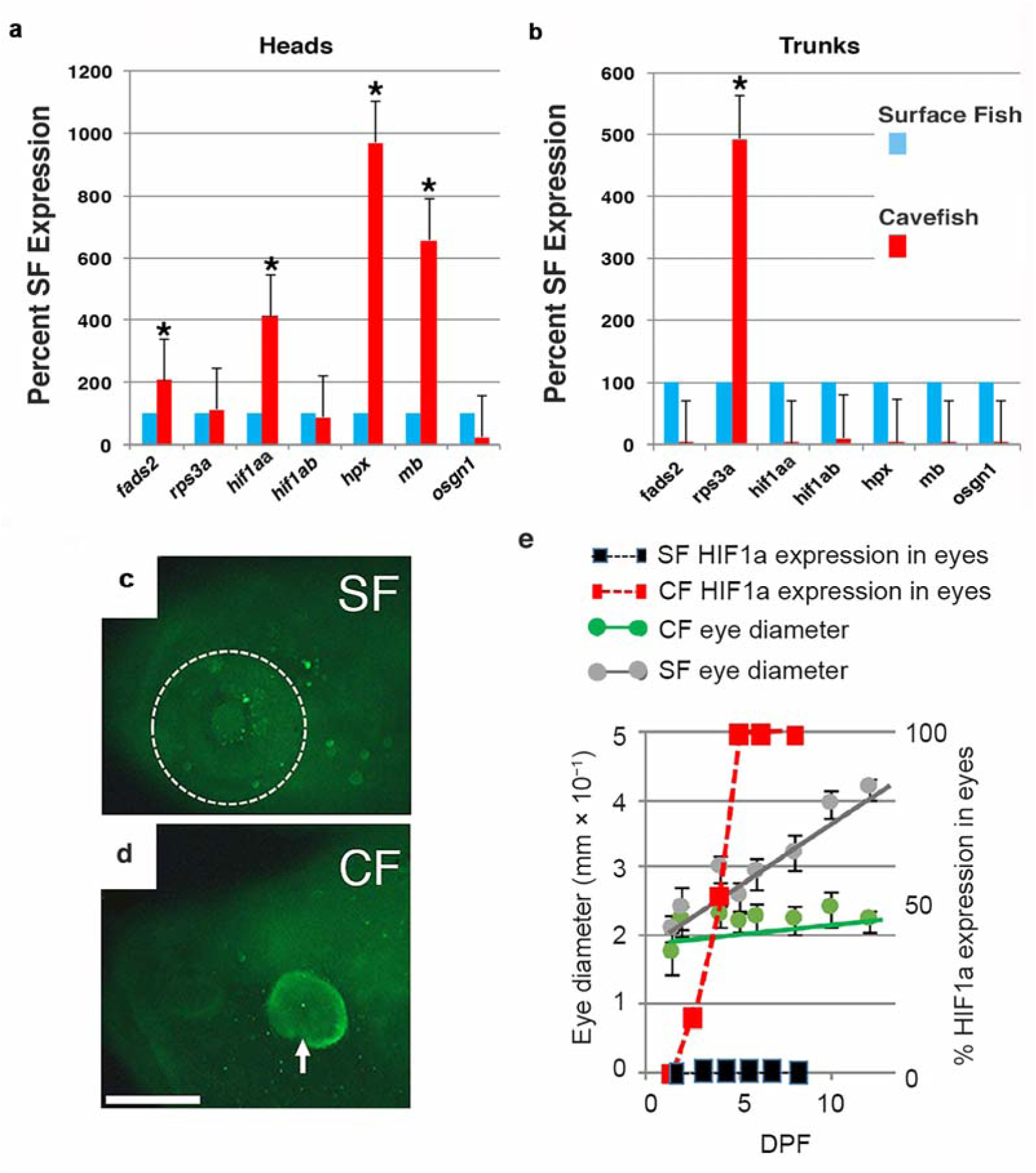
Hypoxia in cavefish. (**a** and **b**). Expression of hypoxia sensitive genes by qPCR in isolated cavefish and surface fish heads and trunks at 1.5 dpf. Histograms show means, and error bars represent s. e. m. of three biological replicates. Asterisks indicate significance at *P* < 0.05 calculated from ΔΔCt values using Student’s two tailed test. (**c** and **d**). Determination of Hypoxia Inducing Factor 1α (HIF1a) expression in SF (c) and PA-CF (d) eyes at 7 dpf by immunostaining. Dotted line indicates SF eye. SF n = 16. PA-CF n = 16. Scale bar: 150 μm; same magnification in K and L. (**e**) HIF1α expression in CF eyes determined by immunostaining compared to optic growth during SF and PA CF development. N for each point = 16.

**Fig. 9.**
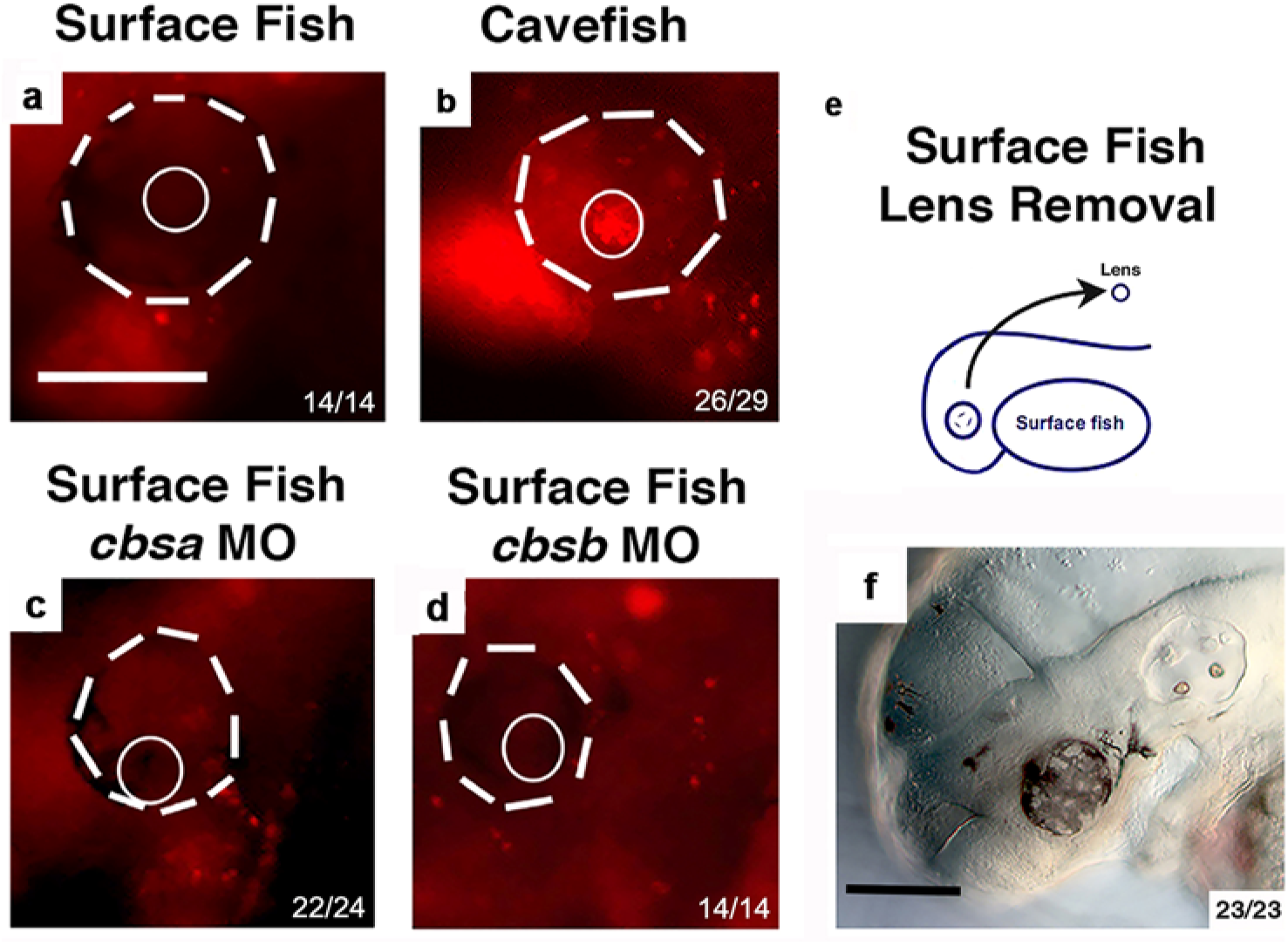
Relationship between *cbs* associated and lens-related optic phenotypes in *Astyanax*. (**a-d**) Lysotracker staining of *cbs* MO injected SF larvae at 68 hpf. (**a**) SF controls showed no lens apoptosis. (**b**) PA-CF controls showed lens apoptosis. (**c** and **d**) *cbsa* morphants (c) and *cbsb* (d) SF morphants showed no lens apoptosis. Dashed lines: eyes. Unbroken lines: lenses. (**e**, **f**) Lens removal from the optic cup on one side of SF larvae at 30 hpf (**e**) did not induce optic hemorrhages at 4 dpf (**f**) or later in development (data not shown). Numbers of specimens with indicated results/total number of analyzed specimens is shown at the lower right side of each frame. Scale bars in A and F: 150 μm; Magnifications are the same in A–L.

### Optic vasculature disruption is independent of lens dysfunction

Previous studies showed that lens apoptosis is an early mediator of CF eye degeneration^9,10^. We conducted the following experiments to understand the relationship between lens dysfunction, *cbsa* downregulation, and optic vasculature defects in CF. First, we injected SF with *cbs* MOs as described above and assayed lens apoptosis in morphant and control larvae at 40 hpf by staining with Lysotracker Red. Apoptotic cells were detected the lens of CF larvae, as described previously^9,10^, but not in SF larvae injected with *cbsa*, *cbsb*, or control MOs, showing that *cbs* knockdown does not promote lens apoptosis (Fig. 8a, d). Second, we removed the lens from the optic cup on one side of a SF head at 30 hpf and determined whether this operation induced optic hemorrhaging later in development. The results showed that optic hemorrhages were absent throughout the remainder of development in SF larvae lacking a lens (Fig. 8f). Together, these results suggest that the lens dysfunction and *cbsa* downregulation/defective optic vasculature development are independent events.

## Discussion

In the present study, we have identified *cbsa* as one of the genes responsible for eye loss in *A. mexicanus* CF. Furthermore, our studies of the hypomorphic *cbsa* phenotype suggest a novel mechanism for CF eye degeneration based on the disruption of optic circulatory function and hypoxia. Our work builds on previous studies carried out during the past several decades that determined the multiplicity of genetic factors involved in CF eye loss^1,2,10^, identified numerous QTL and corresponding genomic regions responsible for the regressive CF eye phenotype^11–13,15^, and tested candidate genes for mutations and roles in CF eye degeneration^17^. However, none of the previous studies were successful in identifying any of the mutated genes responsible for CF eye loss.

Our study, along with information from some of the previous studies mentioned above, provides multiple lines of evidence indicating that *cbsa* is one of the mutated genes responsible for CF eye degeneration. The *cbsa* gene is located near the peak marker of an PA-CF eye size QTL^11,15^, was recognized as a potential candidate for controlling CF eye loss by Ingenuity Pathway Analysis of gene expression data^15^, and is downregulated in eyes and other tissues during larval development in four different CF populations: PA-CF, LS-CF, TI-CF, and MO-CF. Importantly, two or more of these CF populations are thought to have evolved the eyeless phenotype independently, although genetic analysis has shown that they share some of the same genetic factors involved in eye loss^13^. Our results suggest that *cbsa* may be one of the shared genes. Furthermore, we have described a potential *cis*-acting mutation at the same position in the first intron of the *cbsa* gene in six different *A. mexicanus* CF populations (including the four CF populations above), a region that contains an enhancer capable of driving reporter gene expression in SF eyes, brain, and myotomes, the same regions in which *cbsa* is downregulated during CF larval development. We have also presented MO knockdown and CRISPR-Cas9 gene editing results showing that the *cbsa* gene is required for SF eye development. When *cbsa* function was disrupted eye formation was completely abolished or small eyes with ventrally displaced lenses-the typical CF larval eye morphology^2–5^-were formed without affecting general larval morphology. Our results are also consistent with the possibility that the hypomorphic *cbsa* gene is responsible for the phenotypic effects represented by an *Astyanax* eye QTL. However, more comprehensive identification and functional testing of other genes within this QTL interval will be needed to conclude that it is the only gene in the corresponding genomic region that is mutated and involved in eye degeneration. It is also important to note that *cbsa* is one of many genetically^10–13,15^ or epigentically^14^ modified factors that may collaborate to regulate CF eye degeneration, and that other genes remain to be discovered to fully understand this complex phenotype.

Our studies of the CF *cbsa* phenotype revealed that *A. mexicanus* CF show similarities to the human disease homocystinuria and mouse *Cbs* models of this disease with respect to eye abnormalities, elevated hCys levels, and deficiencies in the cardiovascular system^23–27^. We found 4-6 fold increases in hCys levels in PA-CF and TI-CF larvae relative to SF larvae, whereas *Cbs*^+/−^ mice show two-fold elevation of hCys levels, and an approximate 40-fold increase in hCys has been estimated in *Cbs*^−/−^ mice, which die *in utero*^26,27,35^. We showed that *cbsa* expression is decreased in the head of A. *mexicanus* CF larvae, although it remains constant in the trunk, due to strong and probably upregulated expression in the developing CF liver and pancreas. The moderate homocystinura-like effects in CF, and the ability CF to survive them and develop into healthily adults, is likely due to compensation for *cbsa* downregulation by the paralogous *cbsb* gene, which is not downregulated in PA-CF or under a known eye QTL^15^. Consultation of archived ENSEMBL genomic sequence data indicate that duplication of an ancestral *cbs* gene into *cbsa* and *cbsb* paralogs occurred in teleost fish before the split between the families containing zebrafish and *Astyanax*. Zebrafish *cbsa* and *cbsb* show similar patterns of expression and knockout results suggesting that the paralogous *cbsa* and *cbsb* genes show redundant functions^19^. The *A. mexicanus cbsa* and *cbsb* genes are co-expressed in most larval organs, with notable exception of the eyes (and parts of the brain), which is the likely reason that hypomorphic effects of *cbsa* are restricted to the developing CF eyes and do not engender an overall lethal response, such as seen in *Cbs*^−/−^ mice^27^. We also observed *cbsa* expression in the SF brain, as has also been reported for *cbs* genes in zebrafish^19^ and mouse^36^, and *cbsa* downregulation in the CF brain. Brain regions responsible for visual processing, namely the optic tecta, are regressed along with the eyes during CF evolution^37,38^. Therefore, mutated *cbsa* might promote the regression of CF eyes and optic tecta as a functional unit.

Mutations in the *cbs* gene reduce life expectancy in homocystinuria patients because of cardiovascular and other problems and are responsible for cardiovascular defects in mouse CBS models of homocystinuria^24–27^. We discovered defects in CF optic and cranial vasculature, including vascular leakages, eye hemorrhages, and aneurysms, during the critical period of eye degeneration. The localization of these defects to eyes and the brain are attributed to *cbsa* downregulation and the absence of sufficient compensating *cbsb* expression in these areas. In our study, none of the above defects in optic vasculature were seen in more than 1000 SF larvae, except when the *cbs* genes were knocked down by MO injection, implying that *cbs* expression is also required for vasculature development and/or function in *A. mexicanus*. Despite the occurrence of optic vasculature defects, our results have shown that overall CF development is not affected, and survivorship under laboratory conditions is about the same as in SF lacking vascular defects. Thus, the reversibility of homocystinuria-like pathologies in CF highlights the potential of *A. mexicanus* as a natural model for investigating the recovery from hemorrhagic stroke and related cardiovascular diseases.

Our results suggest that inhibition of circulatory system function triggered by *cbsa* downregulation is an important mechanism for inhibiting eye growth in *A. mexicanus*. We substantiated this conclusion by showing that eye growth can be arrested by knockdown of the *rap1b* gene in SF, which prevents vasculature from invading the eyes during larval development. During teleost development, vasculature initially forms in the trunk, and embryonic eye formation in the non-vascularized head is dependent on passive diffusion of oxygen^39^. As larvae increase in size during subsequent development, passive diffusion is no longer sufficient for eye development, and the eyes begin to require oxygen, nutrients, and humoral factors provided by neovascularization in the head^40^. The late reliance of developing eyes on oxygen provided by blood flow would explain why CF form eyes early and eye regression begins later in development. Consistent with this interpretation, our work provides evidence that regressing CF eyes are deficient in oxygen by showing the upregulation of several hypoxia gene markers in the larval head, including the master hypoxia regulator HIF1α, which was restricted to regressing CF eyes. Therefore, we propose a model in which deficiency in oxygen, and/or nutritional and humoral factors, normally provided to growing eyes by the circulatory system, is responsible for optic regression in CF (Fig. 10). Interference with eye growth by blocking the blood supply may confer an evolutionary benefit by eliminating the high energetic cost of maintaining eyes, which are useless in the dark cave environment^41,42^. These findings open the possibility that interference with circulatory system function could also inhibit eye formation in other cave adapted animals, and may have a general role in the evolutionary regression of vestigial organs.

**Fig. 10.**
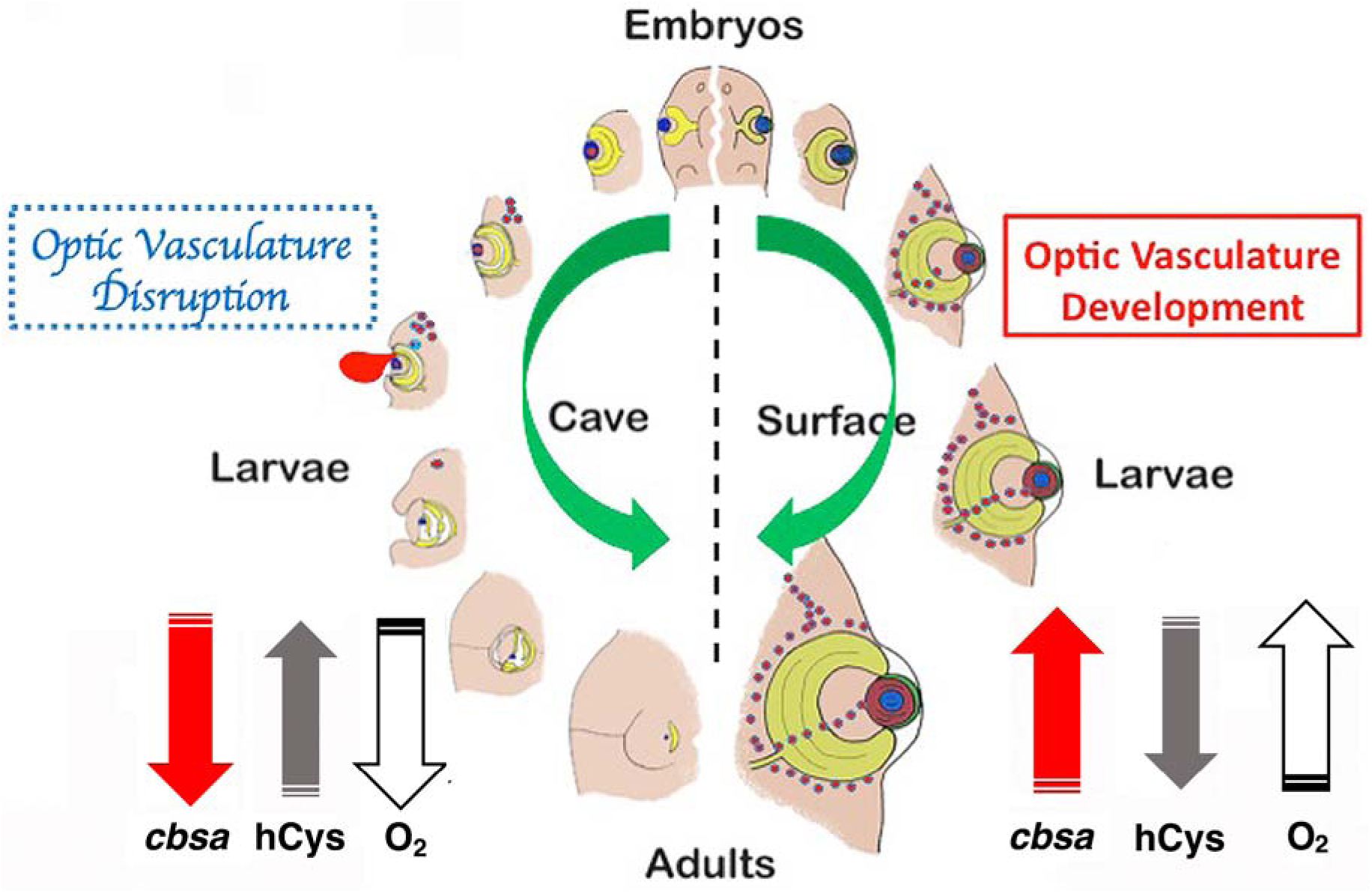
Model for CF eye degeneration based on *cbsa* downregulation, hCys accumulation, and the induction of defects in optic vasculature leading to oxygen deficiency.

Previous studies showed that lens dysfunction is involved in the early stages of CF eye degeneration but these studies also recognized that additional factors must be involved in this process^3,4^. The complexity of CF eye loss was also supported by the multiplicity of genetic factors underlying this process^10–13^. Our results suggest that optic vasculature defects and lens dysfunction are independent processes in the complex CF eye regression phenotype, and that they are likely controlled by different genes. It will be interesting to determine whether other genes underlying this process contribute to the defective optic vasculature phenotype, the lens dysfunction phenotype, both of these phenotypes, or reveal additional changes involved in vestigial eye formation.

## Methods

### Fish Husbandry and Embryo Culture

*A. mexicanus* SF, Pachón (PA-CF), Tinaja (TI-CF), Los Sabinos (LS-CF), Jineo (JI-CF), Chica (CH-CF), and Molino (MO-CF) were obtained from cultured stocks in the Jeffery laboratory. Fish were maintained and spawning was induced as described previously^43^. Embryos and larvae were raised at 23–25°C. Some SF embryos were 400 μM phenylthiourea (Sigma-Aldrich, St. Louis, MO) to reduce pigmentation prior to *in situ* hybridization, blood staining, and immunocytology (see below). Developing SF and PA-CF larvae (40 hpf) were exposed to a 37°C heat shock for 1 hour as described previously^44^. Animals were maintained and handled according to Institutional Animal Care guidelines of the University of Maryland, College Park (IACUC #R-NOV-18-59) (Project 1241065-1).

### Separation of Larval Heads and Trunks

SF and PA larvae were dissected into heads and trunks by making a single incision at the base of the cranium (Fig. 3a) with a fine steel scalpel blade while they were immersed in a small volume of fish culture system water containing 1 μg/ml of MS 222 (Ethyl 3-aminobenzoate methanesulfonate; Western Chemical Inc, Ferndale CA) and viewed under the stereomicroscope. The isolated heads and trunks were immediately immersed in Trizol Reagent (Life Technologies, Grand Island, NY) and RNA was extracted.

### cDNA Synthesis, and RT-PCR

Total RNA was isolated from embryos and larvae using Trizol Reagent, and cDNA was synthesized using the SuperScriptTM III First-Strand Synthesis SuperMix Kit and oligo (dT)_20_ or random hexamer primers (Life Technologies) as described previously^17^. Qualitative RT-PCR was done using the PCR Master Kit (Roche Diagnostics GmbH, Mannheim, Germany). The primers used to amplify protein-coding genes from scaffold KB871589.1 in the Ensembl AstMex1.0.2 PA-CF genome release are listed in Table S3. The primers used for PCR amplification of other genes with expression levels determined by qualitative RT-PCR are listed in Table S4. The PCR cycling conditions for qualitative RT-PCR were 1 cycle of initial denaturation at 94°C for 2 minutes, followed by 25 to 32 cycles of denaturation (94°C for 30 seconds), annealing (60°C for 30 seconds), and elongation (72°C for 30 seconds), and a final elongation step at 72°C for 5 minutes. The standard was 18S rRNA. Quantitative real time RT-PCR (qPCR) of *cbsa*, *cbsb*, and hypoxia sensitive genes was carried out as described by Gore *et al*.^15^ or Ma *et al*.^45^ using GAPDH mRNA as a standard. The primers used for qPCR are listed in Table S5.

### In Situ Hybridization

RNA probes corresponding to gene specific sequences in the *cbsa*, *cryaa*, *hsf2bp*, *cbsb*, *sox2*, *mip*, and *otx2* coding regions were amplified by PCR from SF cDNA using the primers listed in Table S4. The PCR products were cloned into the TOPO vector using the TOPO TA Cloning Kit Dual Promotor (Life Technologies) and confirmed by sequencing. Sense and anti-sense DIG-labeled RNA probes were prepared with SP6 RNA and T7 RNA Polymerases (Roche Diagnostics, Indianapolis, IN). Fixation and processing of samples, whole mount *in situ* hybridization, and photography of hybridized samples was done as described previously^13^.

### F1 Hybrid Test

F1 hybrid embryos were produced by inseminating SF eggs with PA-CF sperm using *in vitro* fertilization procedures^45^. RNA was extracted from isolated heads (see above) of SF, PA-CF, and SF X PA-CF F1 hybrids at 60 hpf. Total RNA was extracted and cDNA was synthesized as described above. The primers for amplification of *cbsa* by RT-PCR were 5’-CGGATGGTGGAAGATGCAGA-3’ (forward) and 5’-CGTAGTGAGCCAGAGGGTTG-3’ (reverse). The PCR cycling conditions were 1 cycle of initial denaturation at 94°C for 2 minutes, followed by 28 cycles of denaturation at 94°C for 30 seconds, annealing at 60°C for 30 seconds, and elongation at 72°C for 30 seconds, and a final elongation step at 72°C for 7 min. Amplification of the control 18S rRNA was carried out using μl of the synthesized cDNA with primers 5’-GAGTATGGTTGCAAAGCTGAAA-3’ (forward) and 5’-CCGGACATCTAAGGGCATCA-3’(reverse). The PCR cycling conditions were 1 cycle of initial denaturation at 94°C for 5 min, followed with 25 cycles of denaturation at 94°C for 30 seconds, annealing at 62°C for 30 seconds, and elongation at 72°C for 30 seconds, and a final elongation step at 72°C for 7 min.

To determine whether the SF or PA-CF *cbsa* allele was expressed in hybrids, PCR reactions were performed to amplify a 327 bp region of the *cbsa* coding region containing the SF- and CF-specific mononucleotide markers using the forward primer 5’-ACCCTCTGGCTCACTACGAC-3’ and the reverse primer 5’-TGCGAGCCATAGCAAAGGAC-3’ and the Phusion High-Fidelity PCR Master Mix (New England Bio Labs, Ipswich, MA) as described previously^17^. The PCR products were purified with the MinElute Gel Extraction Kit (Qiagen, Valencia, CA) and cloned into the Dual Promoter PCR II-TOPO Vector using the TOPO TA Cloning Kit (Life Technologies)^17^. Both PCR products and TOPO plasmids were sequenced. The PCR products were sequenced with forward primer 5’-ACCCTCTGGCTCACTACGAC-3’, and TOPO plasmids were sequenced using the M13-20 and M13rev primers provided with the kit to distinguish which *cbsa* clones contained the SF or CF marker.

### RACE Reactions and Sequencing

The 5’ and 3’ ends of SF and PA *cbsa* mRNA were determined using the SMARTer™ RACE cDNA Amplification kit (Clontech Laboratories, Inc, Mountain View, CA). Poly A^+^ RNA was isolated with the NucleoBond RNA/DNA kit (Macherey-Nagel, Duren, Germany) and RACE-Ready cDNA was generated using the first-strand cDNA synthesis protocol in the SMARTer™ RACE cDNA Amplification kit (Clontech). The primary gene-specific primer for *cbsa* 5’-RACE was 5’-CGCTCTCTCTGCATCTTCCACCATCCG-3’; and the nested gene-specific primer for *cbsa* 5’-RACE was 5’-GCTGATTCTGTCCTTCACACTGCCGCC-3’; the primary gene specific primer for *cbsa* 3’-RACE was 5’-TCTCTGCTCCCCTCACTGTTTTGCCCA-3’; and the nested gene-specific primer for 3’-RACE was 5’-GGAGACGGATCACTTTGCCCTGGTGGT-3’. The PCR reactions were performed using the Advantage 2 PCR Kit (Clontech). The PCR conditions were five cycles at 94°C for 30 seconds and 72°C for 3 minutes; 5 cycles at 94°C for 30 seconds, 70 °C for 30 seconds, and 72°C for 3 minutes, 27 cycles at 94°C for 30 seconds, 68°C for 30 seconds, and 72°C for 3 minutes. For the nested PCR reactions, the cycling conditions were 20 cycles each at 94°C for 30 seconds, 68°C for 30 seconds, and 72°C for 3 minutes. The PCR products were purified with the MinElute PCR Purification Kit (Qiagen) and sequenced with nested gene-specific primers.

### Genome Walking and Sequencing

The SF and PA *cbsa* gene loci and flanking regions were amplified and sequenced by genome walking^17^. Briefly, the entire SF and PA-CF *cbsa* genomic DNA sequences were amplified using the GenomeWalker™ universal kit (Clontech Labotatories, Mountain View, CA) and the overlapping genomic regions were sequenced step by step using the primers listed in Table S6. For the construction of GenomeWalker libraries, genomic DNA was digested with *Eco*RV, *Dra* I, *Pvu*II, and *StuI* I. The GenomeWalker PCR reactions were performed with TaKaRa LA Taq™ (Takara Bio, Mountain View, CA). The PCR reactions were conducted using the two-step cycle parameters described in the GenomeWalker^TM^ kit manual. After obtaining the major bands, the PCR products were inserted into the TOPO vector, and both the PCR products and TOPO plasmids were sequenced.

To amplify SF and PA-CF *cbsa* intron 6, which was not covered by genome walker generated or Ensembl sequences (AstMex1.0.2), we used the forward primer 5’AGATCGTCCGTACCCCTACC-3’, which corresponds to a region located in *cbsa* exon 5, and the reverse primer 5’-TGCCAGCTGAAGTGTGCTTA-3’, which corresponds to a region located in intron 11. The PCR reactions were performed using Phusion High-Fidelity PCR Master Mix (New England Biolabs). The PCR cycling conditions were 1 cycle of initial denaturation at 98°C for 30 seconds, followed by 35 cycles each of denaturation at 98°C for 8 seconds, annealing at 68°C for 25 seconds, extension at 72°C for 5.5 minutes, and a final elongation step at 72°C for 10 minutes. The PCR products were purified with the MinElute Gel Extraction Kit (Qiagen) and sequenced.

To amplify the genomic region between *cbsa* and the adjacent upstream gene *gemin8*, we designed 14 pairs of primers (Table S6) using incomplete sequences available in the Ensembl PA-CF database. The PCR reactions were performed by PCR Master (Roche, Roche Diagnostics GmbH, Mannheim, Germany), and the PCR products were inserted into the TOPO vectors, and both the PCR products and TOPO plasmids were sequenced. After sequencing, low-quality reads and pCRII TOPO vector sequences were removed, the sequences were trimmed, and clean genomic sequence reads were assembled with DNAstar SeqMan Pro software.

### Indel Sequencing in Cavefish Populations

Genomic DNA was extracted from tailfin clips of six different CF populations (see above) using the DNeasy Blood & Tissue Kit (Qiagen). The primers used to amplify the region surrounding the indel in *cbsa* intron 1 were 5’-GCCTGCATGTGCCAGAGGGG-3’ (forward) and 5’-CCGCCGCCAAAACATTGCGT-3’ (reverse). The PCR cycling conditions were one cycle of initial denaturation at 98°C for 30 seconds, followed by 35 cycles of denaturation (98°C for 5 seconds), annealing (at 60°C for 15 seconds), extension (at 72°C for 15 seconds), and a final extension at 72°C for 7 minutes. The primers used to amplify the region surrounding intron 6 were 5’-CCAGAGGCAGACATGTTTCCGATT-3’ (forward) and 5’-GGAGGCTGCAGAGTACTGACAGT-3’ (reverse), and the PCR cycling conditions were the same as those used for the intron 1 region. The primers used to amplify the region surrounding intron 8 were 5’-TGGCTTCAAGCAAGGGCGGG-3’ (forward) and 5’-AGTTGCGGGCAACATCATACCCT-3’ (reverse). The PCR cycling conditions were the same as those used to amplify intron 1, except that annealing was done at 58°C for 15 seconds. All PCR reactions were performed using Phusion High-Fidelity PCR Master Mix (New England Biolabs, Ipswich, MA, USA), and the PCR products were purified with the MinElute PCR Purification Kit (Qiagen) and sequenced.

### Enhancer Transgenesis

Potential enhancers were identified in *cbsa* non-coding DNA sequence using iEnhancer-2L^21^. The predicted enhancer (E in Fig. 1e) in *cbsa* intron 1, which showed major sequence differences between SF and all 6 types of CF, was inserted into the pCMV-GFP vector for expression analysis. The 1140 bp E region was amplified from SF genomic DNA using the forward primer 5’-TGGACGAACGACCAACATTT-3’ and reverse primer 5’-TGCAACTGAAACTGTGCAACT-3’. The pcbsa-CMV-GFP DNA construct containing the predicted enhancer was made by blunt end ligation at the *Spe*I site in the vector. The plasmid was confirmed by sequencing with the EGFP-N primer 5’-CGTCGCCGTCCAGCTCGACCAG-3’. Control (pCMV-GFP) and pcbsa-CMV-GFP DNA constructs and transposase mRNA (25 ng each) were injected into 1-cell SF embryos, and GFP expression was determined by fluorescence microscopy at 40 hpf.

### Gene Knockdown and Rescue

Knockdown of the *cbsa* and *cbsb* genes was done using the splice inhibiting morpholinos *cbsa* MO 5’-CAATGCTAATGCTTTTACCTTCTCC-3’, which targeted *cbsa* exon 5 of the *cbsa* gene, and *cbsb* MO 5’AGCCACTGCAAACACACATACATCA-3’, which targeted *cbsb* exon 4. The *rap1b* translation blocking MO 5’-TCTACAGCCGTGTCTCGTCCTC-3’ targeted part of the 5’ UTR and translation start site of the *rap1b* mRNA. The control MO was 5’-CCTCTTACCTCAGTTACAATTTATA-3’. The MOs were designed and synthesized by Gene Tools, Inc (Philomath, OR). The *cbsa* and *cbsb* MOs were injected into 1–2 cell SF embryos at concentrations of 0.25 mM or 0.5 mM, the *rap1b* MO was injected into 1–2 cell SF embryos at a concentration of 0.125 mM, and the morphants were cultured through the early larval stages. Eye measurements were done in controls and morphants using ImageJ software^45^.

For RT-PCR validations, the *cbsa* and *cbsb* morphants were collected at 40 hpf and total RNA was isolated and cDNA synthesized as described above. The primers used to amplify the region containing the *cbsa* sequence change were 5’-CGGATGGTGGAAGATGCAGA-3’ (forward) and 5’-CGTAGTGAGCCAGAGGGTTG-3’ (reverse). The primers used to amplify the region containing the *cbsb* sequence change were 5’-GGAAAATTGGAGACACGCCG-3’ (forward) and 5’-ATGATGCAGCGGTAACCCTT-3’ (reverse).

For MO rescue, *cbsa* mRNA was microinjected into 1–2 cell *cbsa* or *cbsb* morphant embryos at a concentration of 75 ng/μl in sterile water containing 0.05% phenol red. To prepare *cbsa* mRNA, the full length coding region was amplified from SF cDNA using 5’-GGGCTCGAGCGAATCAGCACCACCTGAAC-3’ (forward) and 5’-GGGTCTAGAAACCATTCCTTTCAGAGACTGGA-3’(reverse) primers, which included *Xho*I and *Xba*I sites. The purified PCR product was digested with *Xho*I and *Xba*I, ligated into the pCS2+ vector, and the chimeric plasmid was confirmed by sequencing. After linearization with *Not*I, the capped mRNA was transcribed using mMESSAGE mMACHINE Sp6 kit (Life Technologies). After transcription, *cbsa* mRNA was recovered by LiCl precipitation, washed in 70% ethanol, re-suspended in sterile H_2_O, and stored at −20°C.

### CRISPR-Cas9 Gene Editing

To knockout the *cbsa* gene in SF, sgRNA and Cas9 mRNA was co-injected into one-cell embryos. The *cbsa* CRISPR targets were designed with the online tool http://crispr.dfci.harvard.edu/SSC/, and the 20-nt target sequence was confirmed online (http://www.e-crisp.org). For sgRNA synthesis, we used a cloning-free, oligo-based method^46^. The sgRNA target sequence in *cbsa* exon 2 was 5’-AGGGTCCATTCTCTTTGCTG-3’, and the sgRNA target sequence in *cbsa* exon 3 was 5’-ATCTTGTTGATGCGCACAAG-3’. The T7 promoter (GG) was added to the 5’ end of the target sequence, and the 20-nt overlapping crRNA-TracrRNA sequence was added to the 3’ end of the sgRNA target sequence. The sgRNA oligo was assembled by PCR using the Phusion High-Fidelity PCR Master Mix (New England Bio Labs, Ipswich, MA). The reaction conditions were denaturation at 98°C for 2 minutes, annealing at 50°C for 10 minutes, and extension at 72°C for 10 minutes. The PCR product was confirmed by agarose gel electrophoresis and purified with the MinElute PCR Purification Kit (Qiagen). The sgRNA was transcribed *in vitro* using the MEGAshortscript High Yield T7 kit (ThermoFisher Scientific) and purified with NucAway^TM^ Spin Columns (ThermoFisher Scientific). For Cas9 mRNA preparation, we used the pST1374-NLS-linker-Cas9 vector (Addgene, USA). The pST1374-NLS-linker-Cas9 (10 µg) was digested with *Pme*I (BioLabs, New England Inc), purified, and 1 ug DNA was used in an *in vitro* transcription reaction with the T7 mMESSAGE mMACHINE kit (ThermoFisher Scientific), and the transcribed RNA was purified using NucAway ^TM^ Spin Columns (ThermoFisher Scientific). We co-injected 25 pg of sgRNA and 300 pg of Cas9 mRNA into one-cell stage SF embryos. CRISPR-Cas9 injected and control un-injected embryos from the same clutch were cultured at 25 °C and phenotyped at 40 hpf by microscopy. Genomic DNA was extracted from embryos with defective eye development and controls, and nested PCR was used to amplify the edited sites. For the exon 2 site, the flanking primers were 5’-GGGAGTCCAAATGAGTGGGT-3’ (forward) and 5’-TTCCCGGGATGGGATTGTGA-3’ (reverse), and the nested primers were -5’-TGGCATACAGCCCTGCTAAA-3’ (forward) and 5’-GTTAGTTAACGGCGCAGACC-3’ (reverse). For the exon 3 site, the flanking primers were 5’-GAGGTCTGCGCCGTTAACTA-3’ (forward) and 5’-AAATCTGTGGCCTGTAGTGCT-3’ (reverse), and the nested primers were 5’-GATGGCTCCTGTGCTGCTAA-3’ (forward) and 5’-AATCTGTGGCCTGTAGTGCT-3’ (reverse). The purified PCR products were assayed for genome targeting efficiency by T7 Endonuclease I digestion and sequenced.

### Homocysteine Determination and Injection

hCys was quantified in SF, PA-CF, and TI-CF larvae using the Homocysteine ELISA Kit (Cell Biolabs Inc, San Diego, CA). Samples containing 100 larvae each were homogenized in PBS, the homogenates were centrifuged at 10,000× *g* for 15 min at 4°C, and the supernatant fraction was collected and assayed immediately according to the instructions of the manufacturer.

L-homocysteine and L-cysteine (both purchased from Sigma Aldrich) were diluted to 1–5 mM in sterile water containing 0.05% phenol red, microinjected into 1–2 cell SF embryos, and the effects on eye development were determined in SF larvae at 4 dpf.

### Microangiography

Microangiography was performed as described previously^47^. Briefly, un-injected SF, PA-CF and *cbsa* or *cbsb* MO injected embryos were mounted in 1.2% low melt agarose prepared in zebrafish embryo medium^48^. Embryos were injected with Qtracker 655 or 705 vascular labels (ThermoFisher Scientific) in the dorsal aorta around the yolk extension area using a pressure injector (World Precision Instruments, Friedberg, Germany). Injected embryos were immediately observed using an upright Leica TCS SP5 confocal microscope. Vascular leakages were analyzed in and around the head regions. At least three to five injected embryos were examined per dose in each experiment.

### Macrophage and Red Blood Cell Staining

Vital staining for macrophages was done by incubating 4–10 dpf PA-CF larvae with 25 μg/ml Neutral Red in fish system tank water for 12 hours at room temperature in the dark^49^. Embryos were stained for red blood cells for 15 minutes with 0.6 mg/ml o-dianisidine (Sigma, St. Louis, MO, USA), 0.01 M sodium acetate (pH 4.5), 0.65% hydrogen peroxide, and 40% (v/v) ethanol in the dark^50^. After staining, the embryos were rinsed in PBS, post-fixed in 4% paraformaldehyde (PFA) for 1 hour, rinsed in PBS, and stored in 80% glycerol.

### Antibody Staining

SF and PA-CF larvae from 1.5–12 dpf were fixed in 4% PFA overnight at 4°C, dehydrated through an increasing series of methanol concentrations to 100% methanol, and stored at −20°C. The larvae were rehydrated, blocked with superblock solution containing 5% goat serum and incubated in a 1:100 dilution of rabbit polyclonal anti-HIF1α primary antibody (Catalogue Number 114977, Abcam, Cambridge, UK) for 48 hours at 4°C, rinsed with superblock, then incubated with 1:200 dilution of Alexa Fluor 488 goat anti-rabbit IgG secondary antibody (ThermoFisher, Waltham, MA) overnight at 4°C, rinsed with PBS, 0.1% Triton X-100, 0.2% BSA, and visualized by fluorescence microscopy. Eye diameters were measured using ImageJ software. Antibody specificity was verified by Western blots.

### Lens Deletion and Apoptosis Detection

The lens was removed from the optic cup on one side of 23 SF embryos at about 30 hpf by microdissection using Tungsten needles^51^ and hemorrhaging was assayed by light microscopy from 2–8 dpf.

Lens apoptosis was determined by vital staining of SF, PA-CF, and *cbs* morphants at 40 hpf with 5 μg/ml Lysotracker Red DND 99 (Invitrogen, Carlsberg, CA) for 30 minutes in the dark^45^. Stained larvae were anesthetized in MS222 (see above) and mounted on glass slides for imaging.

### Statistics

The information used in and provided by statistical test, as well as the statistical analysis used to in these experiments are provided in the figure captions.

### Data Availability

Data are reported in figures, supplementary figures, and supplementary tables, and sequence information has been deposited in NCBI as GenBank accession numbers MK801789 (SF *cbsa* mRNA), MK801790 (PA-CF *cbsa* mRNA), MN186089 (SF *cbsa* genomic region), and MN186090 (PA-CF *cbsa* genomic region).

## Supplementary Information

Supplementary figures

Supplementary tables

## Acknowledgements

We are grateful to Ruby Dessiatoun for fish maintenance and husbandry, to Annie Hasselbalch and Sara Soueidan for technical support, and to Eric Haag, Scott Juntti, and Xiao Liu for comments on a draft manuscript. This work was supported by NIH grant EY024941 to WRJ.

## Author Contributions

L.M., W.R.J., A.V.G., and B.M.W. conceived the project and L.M., A.V.G., W.R.J., D.C., J.S., M.N., and C.M.W performed the research. W.R.J. and L.M. wrote the manuscript with input from all authors.

## References

1. Krishnan, J. & Rohner, N. Cavefish and the basis for eye loss. Philos. Trans. R. Soc. Lond. B Biol. Sci. 372, 20150487 (2017).

2. Jeffery, W. R. Regressive evolution in *Astyanax* cavefish. Annu. Rev. Genet. 43, 25–47 (2009).

3. Yamamoto, Y. & Jeffery, W. R. Central role for the lens in cave fish eye degeneration. Science 289, 631–633 (2000).

4. Strickler, A. G., Yamamoto, Y. & Jeffery, W. R. The lens controls cell survival in the retina: evidence from the blind cavefish astyanax. Dev. Biol. 311, 512–523 (2007).

5. Alunni, A., Menuet, A., Candal, E., Penigault, J-B., Jeffery, W. R. & Rétaux, S. Developmental mechanisms for retinal degeneration in the blind cavefish *Astyanax mexicanus*. J. Comp. Neurol. 505: 221–233 (2007).

6. Fumey, J., Hinaux, H., Noirot, C, Thermes, C., Rétaux, S. & Casane, D. (2018). Evidence for late Pleistocene origin of *Astyanax mexicanus* cavefish. BMC Evol. Biol., 18: 18(1):43. doi: 10.1186/s12862-018-1156-7

7. Herman, A. et al. The role of gene flow in rapid and repeated evolution of cave-related traits in Mexican tetra, *Astyanax mexicanus*. Mol. Ecol. 27, 4397–4416 (2018).

8. Porter, M. L. & Crandall, K. A. Lost along the way: the significance of evolution in reverse. Trends Ecol. Evol. 18: 541–547 (2003)

9. Kim, E. B., et al. Genome sequencing reveals insights into physiology and longevity of the naked mole rat. Nature, 479: 223–227

10. Borowsky, R. & Wilkens, H. Mapping a cave fish genome: polygenic systems and regressive evolution. J. Hered. 93, 19–21 (2002).

11. Protas, M., Conrad, M., Gross, J. B., Tabin, C. & Borowsky, R. Regressive evolution in the Mexican cave tetra, *Astyanax mexicanus*. Curr. Biol. 17, 452–454 (2007).

12. O’Quin, K. E., Yoshizawa, M., Doshi, P. & Jeffery, W. R. Quantitative genetic analysis of retinal degeneration in the blind cavefish *Astyanax mexicanus*. PLoS One 8, e57281 (2013).

13. Borowsky, R. Restoring sight in blind cavefish. Curr. Biol. 18, PR23–R-24 (2008)

14. Gore, A. V. et al. An epigenetic mechanism for cavefish eye degeneration. *Nat*. Ecol. Evol. 2, 1155–1160 (2018).

15. McGaugh, S. E. et al. The cavefish genome reveals candidate genes for eye loss. Nat. Commun. 5, 5307 (2014).

16. Hallmann, K. et al. A homozygous splice-site mutation in CARS2 is associated with progressive myoclonic epilepsy. Neurology 83, 2183–2187 (2014).

17. Ma, L., Parkhurst, A. & Jeffery, W. R. The role of a lens survival pathway including *sox2* and α*A-crystallin* in the evolution of cavefish eye degeneration. EvoDevo 5, 28 (2014).

18. Yoshima, T., Yura, T. & Yanagi, H. Novel testis-specific protein that interacts with heat shock factor 2. Gene 214, 139–146 (1998).

19. Prabhudesai, S. et al. Cystathionine β-synthase is necessary for axis development *in vivo*. Front. Cell Dev. Biol. 6, 14 (2018).

20. Wittkopp, P. J., Haerum, B. K. & Clark, A. G. Evolutionary changes in cis and trans gene regulation. Nature 430, 85–88 (2004).

21. Liu, B., Fang, L., Long, R., Lan, X. & Chou, K. C. iEnhancer-2L: a two-layer predictor for identifying enhancers and their strength by pseudo k-tuple nucleotide composition. Bioinformatics 32, 362–369 (2016).

22. Mudd, S. H., Levy, H. L. & Kraus, J. P. Disorders of transsulfuration. In The Metabolic and Molecular Bases of Inherited Disease (eds C. R. Scriver et al.) 2007–2056 (McGraw-Hill, New York, 2001).

23. Mudd, S. H. et al. The natural history of homocystinuria due to cystathionine β synthase deficiency. Am. J. Hum. Genet. 37, 1–31 (1985).

24. Kraus, J. P. & Kožich, V. Cystathionine-β Homocysteine in Health and Disease (eds R. Carmel & D. W. Jacobsen) 223–243 (Cambridge University Press, Cambridge, 2001).

25. Cai, Y. et al. Homocysteine-responsive ATF3 gene expression in human vascular endothelial cells: activation of c-Jun NH_2_-terminal kinase and promoter response element. Blood 96, 2140–2148 (2000).

26. Kamath, A. F. et al. Elevated levels of homocysteine compromise blood-brain barrier integrity in mice. Blood 107, 591–593 (2006).

27. Tawfik, A. et al. Alterations of retinal vasculature in cystathionine-beta-synthase mutant mice, a model of hyperhomocysteinemia. Inves. Ophthalmol. Vis. Sci. 54, 939–949 (2013).

28. Gore, A. V., Lampugnani, M. G., Dye, L., Dejana, E. & Weinstein, B. M. Combinatorial interaction between CCM pathway genes precipitates hemorrhagic stroke. Dis. Model. Mech. 1, 275–281 (2008).

29. Long, Y. et al. Transcriptional events co-regulated by hypoxia and cold stresses in zebrafish larvae. BMC Genomics 16, 385 (2015).

30. Ton, C., Stamatiou, D. & Liew, C. C. Gene expression profile of zebrafish exposed to hypoxia during development. Physiol. Genomics 13, 97–106 (2003).

31. Alvarez-Tejado, M. et al. Hypoxia induces the activation of the phosphatidylinositol 3-kinase/Akt cell survival pathway in PC12 cells: protective role in apoptosis. J. Biol. Chem. 276, 22368–22374 (2001).

32. Deuel, J. W. et al. Different target specificities of haptoglobin and hemopexin define a sequential protection system against vascular hemoglobin toxicity. Free Radic. Biol. Med. 89, 931–943 (2015).

33. Wittenberg, B. A. & Wittenberg, J. B. Transport of oxygen in muscle. Ann. Rev. Physiol. 51, 857–878 (1989).

34. Mazurais, D. et al. Identification of hypoxia-regulated genes in the liver of common sole (*Solea solea*) fed different dietary lipid contents. Mar. Biotechnol. (NY) 16, 277–288 (2014).

35. Watanabe, M., et al. Mice deficient in cystathionine ß-synthase: animal models for mild and severe homocyst(e)inemia. Proc. Natl. Acad. Sci USA. 92, 1585–1589. (1995)

36. Enokido, Y. et al. Cystathionine ß-synthase, a key enzyme for homocysteine metaboloism, is preferentially expressed in the radial glia/astrocyte lineage of the developing mouse CNS. FASEB J. 10.1096/fj.05-3724fje. (2016)

37. Schmatolla, E. Dependence of tectal neural differentiation on optic innervation in teleost fish. J. Embryl. exp. Morph. 27, 555–576. (1972)

38. Soares, D. Yamamoto, Y., Strickler, A. G, & Jeffery, W. R. The lens has a specific influence of optic nerve and tectum development in the blind cavefish *Astyanax*. Dev. Neurosci, 26, 308–317 (2004)

39. Luo, X., Shen, Y. M., Jiang, M. N., Lou, X. F. & Shen, Y. Ocular blood flow autoregulation mechanisms and methods. J. Ophthalmol. 2015, 864871 (2015).

40. Pelster, B. & Burggren, W. W. Disruption of hemoglobin oxygen transport does not impact oxygen-dependent physiological processes in developing embryos of zebra fish (*Danio rerio*). Circ. Res. 79, 358–362 (1996).

41. Yu, D. Y. & Cringle, S. J. Oxygen distribution and consumption within the retina in vascularised and avascular retinas and in animal models of retinal disease. Prog. Retin. Eye Res. 20, 175–208 (2001).

42. Moran, D., Softley, R. & Warrant, E. J. The energetic cost of vision and the evolution of eyeless Mexican cavefish. Sci. Adv. 1, e1500363 (2015).

43. Jeffery, W. R., Strickler, A. G., Guiney, S., Heyser, D. G. & Tomarev, S. I. Prox 1 in eye degeneration and sensory organ compensation during development and evolution of the cavefish astyanax. Dev. Genes Evol. 210, 223–230 (2000).

44. Hooven, T. A., Yamamoto, Y. & Jeffery, W. R. Blind cavefish and heat shock protein chaperones: a novel role for hsp90alpha in lens apoptosis. Int. J. Dev. Biol. 48, 731–738 (2004).

45. Ma, L. et al. Maternal genetic effects in *Astyanax* cavefish development. Dev. Biol. 441, 209–220 (2018).

46. Varshney, G.K. et al. A high-throughput functional genomics workflow based on CRISPR/Cas9-mediated targeted mutagenesis in zebrafish. Nat. Protocols 11, 2357–2375 (2016).

47. Jung, H. M. et al. Imaging blood vessels and lymphatic vessels in the zebrafish. Methods Cell Biol. 133, 69–103 (2016).

48. Westerfield, M. The Zebrafish Book: A Guide for the Laboratory Use of Zebrafish (Danio rerio). 5 edn, (University of Oregon Press, Eugene, OR, 2007).

49. Herbomel, P., Thisse, B. & Thisse, C. Zebrafish early macrophages colonize cephalic mesenchyme and developing brain, retina, and epidermis through a M-CSF receptor-dependent invasive process. Dev. Biol. 238, 274–288 (2001).

50. O’Brien, B. R. A. Identification of haemoglobin by its catalase reaction with peroxide and o-dianisidine. Stain Technol. 36, 57–61 (1961).

51. Yamamoto, Y. & Jeffery, W. R. Probing teleost eye development by lens transplantation. Methods 28, 420–426 (2002)

