## Supplementary Figures and Tables for "A Hypomorphic *Cystathionine ß-Synthase* Gene Contributes to Cavefish Eye Loss by Disrupting Optic Vasculature"

### Supplementary Figures and Legends

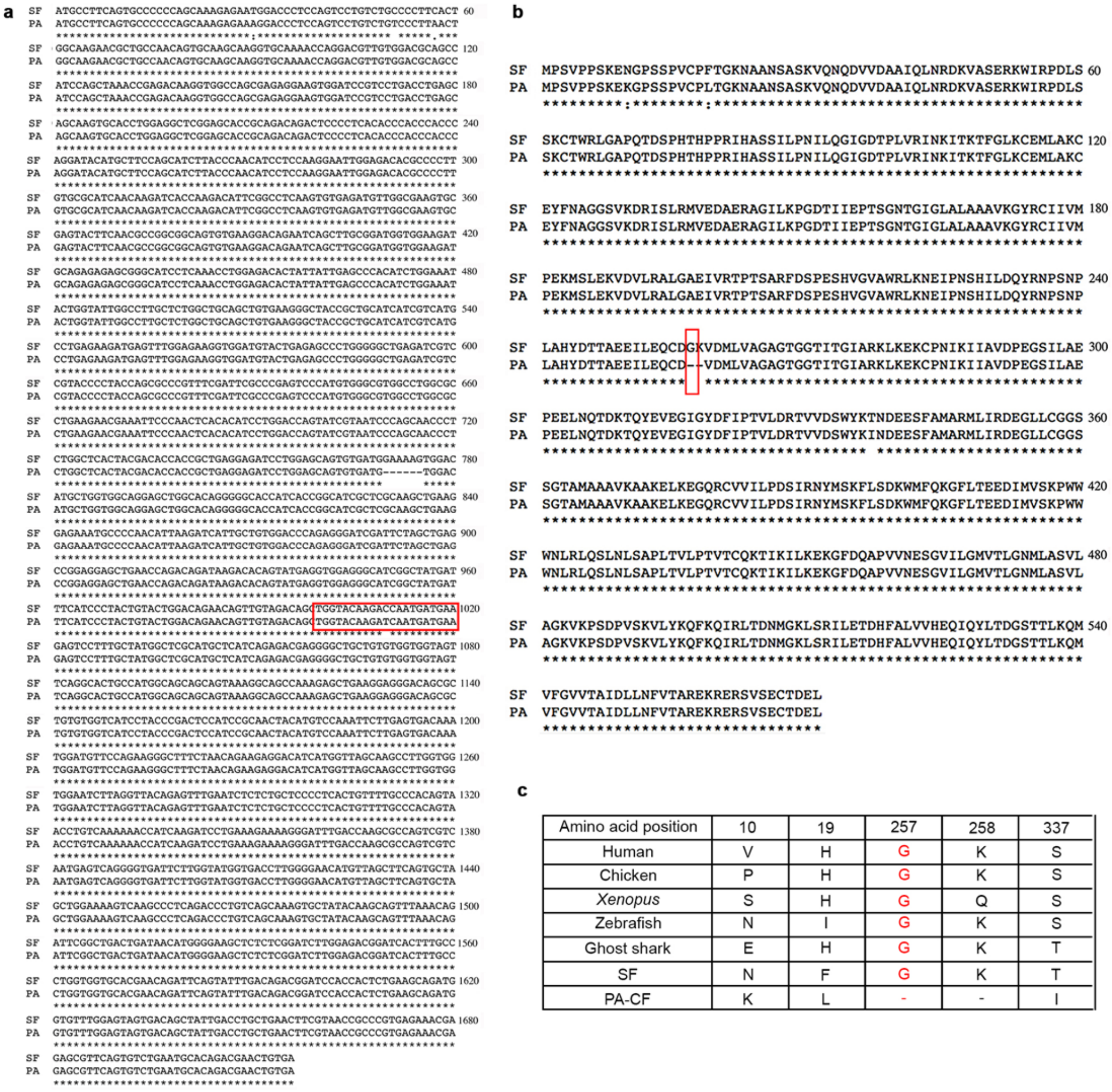

| Amino acid position | 10 | 19 | 257 | 258 | 337 |
|---|---|---|---|---|---|
| Human | V | H | G | K | S |
| Chicken | P | H | G | K | S |
| *Xenopus* | S | H | G | Q | S |
| Zebrafish | N | I | G | K | S |
| Ghost shark | E | H | G | K | T |
| SF | N | F | G | K | T |
| PA-CF | K | L | - | - | I |

**Supplementary Fig. 1: Nucleotide and protein sequence of the SF and PA-CF *cbsa* genes**. (**a**) Nucleotide coding sequence alignments the SF and PA-CF *cbsa* genes. Asterisks: identical nucleotides. Spaces, :, or - - - show different nucleotides or deletions respectively. The region complementary to the sequence used to identify the SF and PA-CF alleles in the F1 hybrid test, including the SNP marker (Fig. 3a), is boxed in red. (**b**) Deduced amino acid sequence alignment of the SF and PA-CF CBSA proteins.

Asterisks: identical amino acids. · or - - -: different amino acids or deletions respectively. The amino acid that is conserved in vertebrates and SF but absent in PA-CF CBSA (G-257) is boxed in red. (**c**) Comparison of divergent amino acid sites in SF and PA-CF CBSA to conserved amino acids at the same position in CBSA (zebrafish) or other vertebrate CBS proteins. Variable amino acid sites are indicated in black type. The conserved amino acid site (G-257) is indicated in red type. Dashes indicate missing amino acids in PA-CF CBSA.

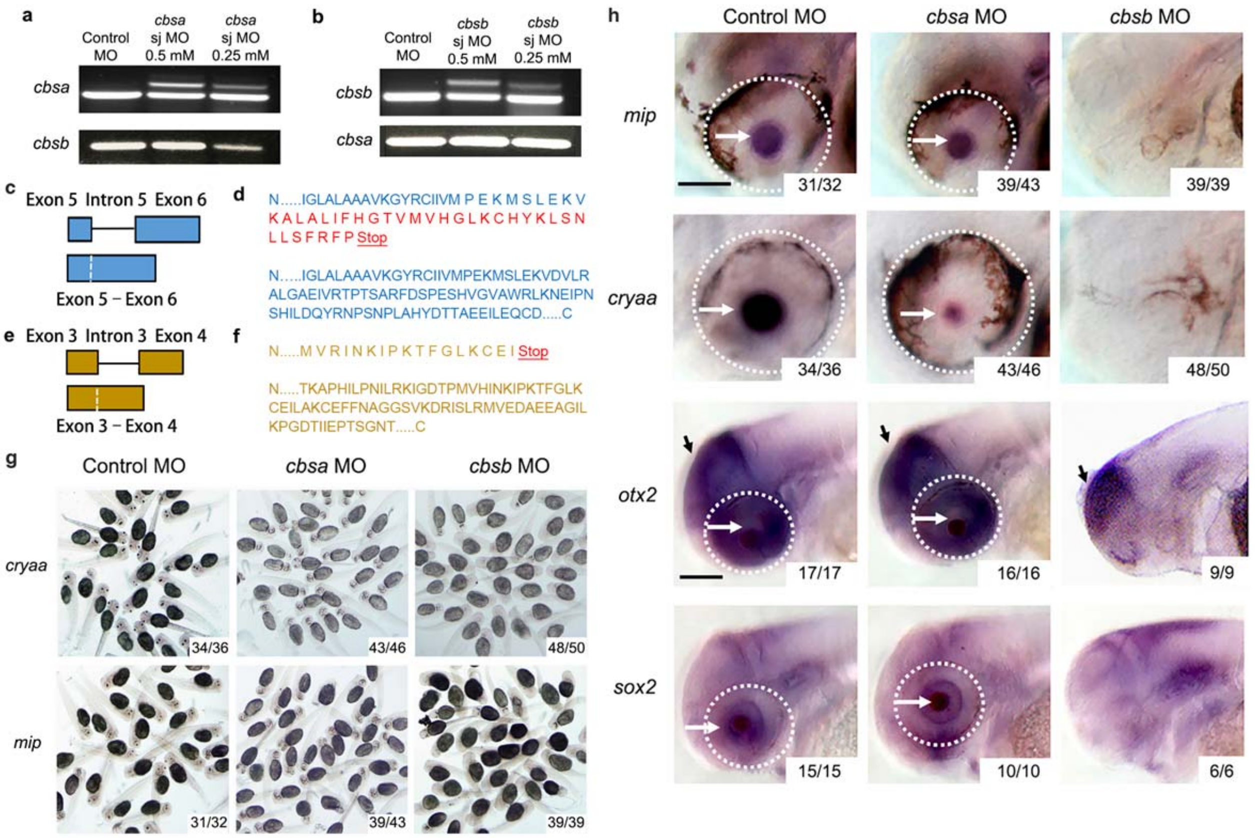


**Supplementary Fig. 2: Validations (a-e) and controls ( g-h) for morpholino based *cbs* gene knockdown**. (**a-b**). Validations by RT-PCR showing *cbsa* and *cbsb* expression in morphants produced by injection of 0.5 mM or 0.25 mM *cbsa* (top) or *cbsb* (bottom) MOs into SF eggs. The upper bands in the *cbsa* and *cbsb* MO lanes are PCR products from pre-mRNAs containing unprocessed *cbsa* or *cbsa* introns including translation stop sites (see d below). The lower bands are PCR products from processed mRNAs. (**e**) Diagrams of the *cbsa* (top, blue) and *cbsb* (bottom, yellow) intron-exon boundary regions (top) and processed exons (bottom) targeted by *cbsa* and *cbsb* splice junction MOs. (**d**) Deduced amino acid sequences of *cbsa* of sequenced upper and lower PCR products in I. The deduced amino acid sequences of PCR products from the *cbsa* and *cbsb* upper transcripts both contain unprocessed stop codons in introns that would lead to premature translation termination. *cbsa* exons: blue letters. *cbsb* exons: yellow letters. Introns and translation termination (stop) sites: red letters. (g-h). Controls. *In situ* hybridized groups of 40 dpf larvae showing (**g**) *cryaa* and *mip* expression in SF injected with 0.5 mM control, *cbsa* or *cbsb* MO. (**h**) Eye morphology and *in situ* hybridization showing *mip*, *cryaa*, *otx2*, and *sox2* expression in SF injected with 0.5 mM control MO, *cbsa* MO, or *cbsb* MO. Number of larvae showing the indicated results/ total number of specimens is shown at the lower right in A. Dashed lines: eyes (when present). White arrows: lens. Black arrows: forebrain. Scale bars: 150 μm; magnification is the same in all frames in the top and the bottom two rows.

**Supplementary Tables**

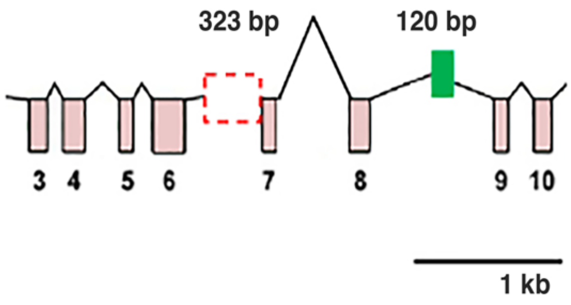


| Cavefish Type | Intron 6<br>323 bp<br>deletion<br>indel | Intron 8<br>120 bp insertion<br>indel |
|---|---|---|
| PA-CF | + | + |
| TI-CF | + | + |
| LS-CF | + | + |
| CH-CF | + | + |
| ML-CF | - | - |
| JI-CF | - | - |

**Supplementary Table 1**. The presence (+) or absence (-) of indels in *cbsa* introns 6 and 8 in different CF populations compared to SF.

**a**

| Egg injection | Total number | Number of normal eyes | Number of small or absent eyes | Percent of effected eyes (%) |
|---|---|---|---|---|
| Cys | 122 | 114 | 8 | 6.5 |
| hCys | 133 | 104 | 29 | 21.8 |

**b**

| Morpholino | Total number | Number of normal eyes | Number of small or absent eyes | Percent of effected eyes (%) |
|---|---|---|---|---|
| Control | 450 | 437 | 18 | 4.0 |
| *cbsa* | 511 | 306 | 195 | 38.1 |
| *cbsb* | 408 | 16 | 364 | 89.2 |

**c**

| Injection | Total number | Number of large eyes | Percent of large eyes (%) | Number of small eyes with ventrally displaced lens | Percent of small eyes (%) | Number of absent eyes | Percent of absent eyes (%) |
|---|---|---|---|---|---|---|---|
| *cbsa* MO | 28 | 8 | 28.6 | 16 | 57.1 | 4 | 4.3 |
| *Cbsa* MO + *cbsa* mRNA | 40 | 34 | 85 | 4 | 10 | 2 | 5 |
| *cbsb* MO | 48 | 2 | 4.2 | 10 | 20.8 | 36 | 75 |
| *cbsb* MO + *cbsa* mRNA | 45 | 0 | 0 | 32 | 71.1 | 13 | 28.9 |

**Supplementary Table 2**. (**a**) Effect of 1 mM homocysteine (hCys) or cysteine (Cys) injection into eggs on SF larval eye development at 40 hpf. (**b**) Effects of splice junction (sj) *cbsa* and *cbsb* morpholinos on eye development in SF at 40 hpf. **(c)** Rescue of eye development after morpholino (MO) knockdown by mRNA injection.

**Supplementary Table 3.** Gene models and ID numbers of protein coding genes on scaffold KB871589.1 in the Ensembl AstMex102 genome assembly and primers used for screening for gene expression levels in SF and PA at 40 hpf by qualitative RT-PCR.

| Gene name | Ensembl version gene ID | Forward primer (5’-3’) | Reverse primer (5’-3’) |
|---|---|---|---|
| *arhgap6-2* | ENSAMXG00000007866 | TCACCATCCCCAAAGATGGC | GGAGTCTTTGTGCTCCTCCC |
| *arhgap6-3* | ENSAMXG00000007844 | TTCATGACGTGGCTGCGTT- | AGATCCTCGAACGTGGCGAT |
| *msl3-201* | ENSAMXG00000007892 | GTTCCCCACTGTTTTGAGCG | TTGCGTGCCAGTTTACGTTG |
| *frmpd4* | ENSAMXG00000007924 | GACTCACCAAGGAGTGCGAA | GGAGCCAGAGAGGGTAAGGA |
| *pwp2h* | ENSAMXG00000007988 | CCTGTTCACAGCGTCTCCTT | TCCGTCAGGAGAATAGGCCA |
| *tmsb4x* | ENSAMXG00000008001 | GAGGTCACCAGCTTCGACAA | ATGACGCTTGCTTCTCCTGT |
| *tceanc* | ENSAMXG00000025888 | AGCTGCAGACCACTGACATC | CACAGACATGCGGGCAAAAA |
| *rab9a* | ENSAMXG00000025889 | GGACGGCTGGTCACTTTACA | CCACCTTTGCGCTTCCTCTA |
| *gpm6ba* | ENSAMXG00000008016 | ATGTGGGCAAAACTTGGCTG | GCGTCGTCCTTGGTCTTGAT |
| *gemin8* | ENSAMXG00000008029 | CTGTCTATGCCCGCTACTGG | GTCTGTGCAAAGAACTGCCG |
| *cbsa* | ENSAMXG00000008046 | CGCATGCTCATCAGAGACGA | GGCAAAGTGATCCGTCTCCA |

| | | | |
|---|---|---|---|
| *cryaa* | ENSAMXG00000008095 | TTTGACTATGACCTCTTCCCCTACGC | GGGGGTAGAGTTAGTCTTGTCGTCAC |
| *hsf2bp* | ENSAMXG00000008124 | AGGAGCAGAAGAAGCAGCAG | ATAAGCGGTATGAGTGCGGG |
| *ankrd10* | ENSAMXG00000008136 | AACGCAAATGGGTTGACTGC | TCTCCGCATGATCAGGTTCG |
| *ing1* | ENSAMXG00000008150 | CCTGAGGGGACTCCCTTTGA | AGCTGGACTGTTTGGGTTCG |
| *cars2* | ENSAMXG00000008172 | CCTGGTACAGTTGTGGACCC | CCCCATGGAGACTCCCAGTA |
| *rab20* | ENSAMXG00000008185 | CCTGCGCCCAGAAAAATGAG | ATGGAACTGCTCACGACCTG |
| *irs2* | ENSAMXG00000025890 | GCTCTGTCTTAGGATCGCCC | CCTTGGTGTTTGTTGGAGCG |
| *lig4* | ENSAMXG00000008192 | TCGCCCTCACAGCGTATTTT | CGTTGAGGGCGTTGACTACT |
| *fam155a* | ENSAMXG00000008195 | TATACACGGAGGCACTCCCA | GGGAGATGGGTTTTGGAGCA |

**Supplementary Table 4**. Oligonucleotide primers used to prepare probes for *in situ* hybridization or to amplify *atf3* and *shha* RNA by qualitative RT-PCR.

| Transcript | Forward primer (5’-3’) | Reverse primer (5’-3’) |
|---|---|---|
| *cbsa* (ISH) | CGCATGCTCATCAGAGACGA | GGCAAAGTGATCCGTCTCCA |

| | | |
|---|---|---|
| *hsf2bp* (ISH) | AGGAGCAGAAGAAGCAGCAG | ATAAGCGGTATGAGTGCGGG |
| *cbsb* (ISH) | CGATGGTGCGCATCAACAAA | TGACGATGATGCAGCGGTAA |
| *sox2* (ISH) | CTGCACATGAAGGAACACCC | GACATGCTGTAGGTGGGCGA |
| *mip* (ISH) | ACTTTTGCCTTCCTGATCG | AGGTGTCCCATGAGCACAGA |
| *otx2* (ISH) | ATGATGTCGTATCTCAAGCAACC | TAATCCAAGCAGTCAGCATTGAAG |
| *cryaa* (ISH) | TTTGACTATGACCTCTTCCCCTACGC | GGGGGTAGAGTTAGTCTTGTCGTCAC |
| *shha* (RT-PCR) | TATGAAGGCCGGGCCGTGGA | CCGGGTACGACGTTGCTCGC |
| *atf3* (RT-PCR) | ACGCTCGACGACTTTACCAC | GGTGCTCTCCTTGATCTGCT |

**Supplementary Table 5**. Oligonucleotide primers used in quantitative real-time RT-PCR determinations.

| Gene name | Ensembl version gene ID | Forward primer (5’-3’) | Reverse primer (5’-3’) |
|---|---|---|---|

| | | | |
|---|---|---|---|
| *fads2* | ENSAMXG00000015974 | TGCAACGGAAGCGT TTACAG | CTGGAAATCTGCAA CCAGGG |
| *rps3a* | ENSAMXG00000021691 | TGTTCAACATCCGCA ACCTG | CGGAGGCGATTTTA GTTCCC |
| *hif1αa* | ENSAMXG00000039550 | GCACTTTACCTACTG CGACG | TGGCAAACAAGTTG TGGTGA |
| *hif1αb* | ENSAMXG00000019342 | TACAACAGGGATGT CTGCGG | TGGGCATTCTGGATG GCTAA |
| *hpx* | ENSAMXG00000002129 | TATGCTTTCAGAGGC CACCA | TAGGAGAAGACAGC GTCCAC |
| *mb* | ENSAMXG00000030396 | AAAGTTGGGAATCG GTCGGA | CCATGGCCTCGGAT GAGTT |
| *osgn1* | ENSAMXG00000037872 | CCGAACCCGAACAC CCA | TCCATGACTGAAGCT CGGG |
| *cbsa* | ENSAMXG00000008046 | ACAGATTCGGCTGAC TGATAAC | GAGAAGAGGTGTGC TCAAACT |
| *cbsb* | ENSAMXG00000018461 | CTGGAGCAGTGTGAT GGTAAA | GGGTCCACTCCAAC AATCTT |
| *GAPDH* | ENSAMXG00000039361 | TCCTGAACTCAATGG CAAGC | TTCTCCAAGCGGAC AGTCAA |

**Supplementary Table 6**. Primers used for PCR amplification and genome walking in the SF and PA *cbsa* gene loci.

| Primer | Primer sequence 5’-3’ |
|---|---|
| Ma171 | GTTATGCCTGAGAAAATGAG- |

| | |
|---|---|
| Ma172 | AAGAACTTAGACATGTAGTT |
| Ma173 | AAAGTGGATATGCTGGTGGC |
| Ma174 | AAGTTGAGAAGATCAATGGC |
| Ma175 | ACGGATCCACCACTCTGAAGCA |
| Ma176 | GCTCTCGTTTCTCACGGGCGG |
| Ma177 | TTTGACAGACGGATCCACCACTCT |
| Ma178 | ACGCTCTCGTTTCTCACGGGC |
| Ma305 | GAGCAGTGTGATGGTTGTGG |
| Ma306 | CGGGTAGGATGACCACACAG |
| Ma307 | CGCATGCTCATCAGAGACGA |
| Ma308 | GGCAAAGTGATCCGTCTCCA |
| Ma329 | CGCTCTCTCTGCATCTTCCACCATCCG |
| Ma330 | GCTGATTCTGTCCTTCACACTGCCGCC |
| Ma331 | TCTCTGCTCCCCTCACTGTTTTGCCCA |
| Ma332 | GGAGACGGATCACTTTGCCCTGGTGGT |
| Ma339 | TCACCAAGACATTCGGCCTC |
| Ma340 | AAGCAGTGGTATCAACGCAG |
| Ma341 | TCGTTGTAAATCGACGGCCA |
| Ma342 | CACAGTTGGTGTTGTTTTTGAGC |
| Ma343 | CAGCCACCCAGATAAAGCGA |
| Ma344 | AAGCAGTGGTATCAACGCAGA |
| Ma345 | TGGGAGAAGAGAAAACAGCGG |
| Ma346 | ACAGTTGGTGTTGTTTTTGAGCTT |
| Ma347 | CGGCAGTGTGAAGGACAGAA |
| Ma348 | GAACGCTCTCGTTTCTCACG |
| Ma349 | AGTGCGAGTACTTCAACGCC |
| Ma350 | AGACACTGAACGCTCTCGTTT |
| Ma351 | ACGGCCATGTGTGTTTGTTG |
| Ma352 | GAGGCCGAATGTCTTGGTGA |
| Ma353 | CATAATGGCCTGGGGGATAGC |
| Ma354 | GAAGTACTCGCACTTCGCCA |
| Ma405 | TGTCAGTGAACAATGTTGATGTT |
| Ma406 | GGATCAAATACTTCTTGGACTCAC |
| Ma407 | CAGACAGTAGTGTACAGATTCCCA |
| Ma408 | CACCGGTTGTTCAGGTCCAT |
| Ma409 | AGTAGTGTACAGATTCCCATTGC |
| Ma410 | TGCATGGATGGGGTGTTTGT |
| Ma437 | CGGATGGTGGAAGATGCAGA |
| Ma438 | CGTAGTGAGCCAGAGGGTTG |
| Ma439 | CAGAATCAGCTTGCGGATGG |
| Ma440 | CTGGTAGGGGTACGGACGAT |
| Ma441 | CGATGGTGCGCATCAACAAA |
| Ma442 | TGACGATGATGCAGCGGTAA |
| Ma443 | GGAAAATTGGAGACACGCCG |
| Ma444 | ATGATGCAGCGGTAACCCTT |

| | |
|---|---|
| Ma445 | AATCAGCACCACCTGAACCT |
| Ma446 | CAGCCACCGCGAAAATGAC |
| Ma447 | GGATATGCTGCAAGCCGAGA |
| Ma448 | CGGGGTAAAGTTCACGGGAG |
| Ma449 | GCCGAGAAAATGGGTAAAGCC |
| Ma450 | TCCGTTATCGGGGAAGAAGG |
| Ma451 | TCACCAAGACATTCGGCCTC |
| Ma452 | TGTAAAACGACGGCCAGTGA |
| Ma453 | CAGCCACCCAGATAAAGCGA |
| Ma454 | TTGGGTAACGCCAGGGTTTT |
| Ma481 | TGGGTGGGTGGGTGTGAGGGGAGTCTGT |
| Ma482 | GGTGCTCCGAGCCTCCAGGTGCACTT |
| Ma483 | TCTCGCTGGCCACCTTGTCTCGGTTTA |
| Ma484 | TCCTGTCTGCCCCTTCACTGGCAAGAA |
| Ma485 | CTGCCAACAGTGCAAGCAAGGTGCAAAA |
| Ma486 | CGTTGTGGACGCAGCCATCCAGCTAAA |
| Ma487 | TCACCAAGACATTCGGCCTC |
| Ma488 | CCATGGCAGTGCCTGAACTA |
| Ma489 | AGATCGTCCGTACCCCTACC |
| Ma490 | TGCCAGCTGAAGTGTGCTTA |
| Ma491 | TGGGGTTTGGTGCTCGTATG |
| Ma492 | AGACAGCCTTGCACCTACAG |
| Ma 493 | GGTCCTAAAGCCTGTTGGGT |
| Ma 494 | TGTTGGTCGTTCGTCCAATTAT |
| Ma 495 | TGGACGAACGACCAACATTT |
| Ma 496 | TGGCACAACCCATATTTGCAT |
| Ma 497 | AATTGGACGAACGACCAACA |
| Ma 498 | CTTAGCTGCATATCATTGAA |
| Ma 499 | AGAACTCTGACTCAGTTGTACATT |
| Ma 500 | AGACACTTAATAGGACCACAAACA |
| Ma 501 | GCCAGTCTGATCTGGTGCTT |
| Ma 502 | TGGCGGTTACTTATCATGTGT |
| Ma 503 | TGTAAGAAGAGCAACATTACGGT |
| Ma 504 | ATCTATCCTGCCCCTTGCAC |
| Ma 505 | GCCTGCTGGACCACTCAAA |
| Ma 506 | GGATCTGAATGATGTCTGTT |
| Ma 507 | GGTGAAGTGATGGACCTGTT |
| Ma 508 | GGACTATACCTATTATACT |
| Ma 509 | GGGAGCTTTTTAACTGTCAG |
| Ma 510 | CTACTGAACATTAGTGGTGCA |
| Ma 511 | ACTGTTCTGTCCAGTACAG |
| Ma 512 | CGATCAGCATCACTTTACTT |
| Ma 513 | CCTGTTAGCCCATATCAGAT |
| Ma 514 | TGACGCACTGATCTACGTGA |
| Ma 515 | CCTCGATTAGCATATTGTCA |

| | |
|---|---|
| Ma516 | TGCCGCCGGCGTTGAAGTACTCGCACTT |
| Ma517 | GTTGATGCGCACAAGGGGCGTGTCTCCAA |
| Ma518 | GGCACGTGGGGTTTGGTGCTCGTATGT |
| Ma519 | ACAGCGCGTTCCCACACGTGGCTCGTT |
| Ma520 | TGGCTGCCTTTACTGCTGCTGCCATGGC |
| Ma521 | ACCACACAGCAGCCCCTCGTCTCTGATG |
| Ma522 | GGGTGATTCTTGGTATGGTGACCTTGG |
| Ma523 | ATGGGGAAGCTCTCTCGGATCTTGGAG |
| Ma524 | TGCTCAAGTGTGCTTGTCAG |
